# Involvement of OsS40-14 in ROS and plastid organization related regulatory networks of dark-induced leaf senescence in rice

**DOI:** 10.1101/2024.08.01.606232

**Authors:** Habiba, Chunlan Fan, Wuqiang Hong, Ximiao Shi, Xiaowei Wang, Weiqi Wang, Wenfang Lin, Yanyun Li, Noor ul Ain, Ying Miao, Xiangzi Zheng

**Affiliations:** Fujian Provincial Key Laboratory of Plant Functional Biology, Fujian Agriculture and Forestry University, 350002 Fuzhou, China

**Keywords:** OsS40, dark-induced rice leaf senescence, core binding motif, integrative analysis of meta-datasets, plastid organization

## Abstract

Dark-induced senescence triggers significant metabolic changes that recycle resources and ensure plant survival. In this study, we identified a transcription factor OsS40-14 in rice, which can form homo-oligomers. The *oss40-14* knockout mutants exhibited stay-green phenotype of primary leaf and flag leaf during dark-induced condition, with substantial retention of chlorophylls and photosynthetic capacity as well as remarkably reduced reactive oxygen species (ROS), while *OsS40-14* overexpressing transgenic lines (*oeOsS40-14*) showed an accelerated senescence phenotype under dark-induced leaf senescence conditions. Transcriptome analysis revealed that when the detached leaves of *oss40-14* and WT were treated in darkness condition for 72 hours, 1585 DEGs (|Log2FC| ≥1, P value<0.05) were reprogrammed in *oss40-14* relative to WT. CUT&Tag-seq analysis in protoplast transient expression of OsS40-14 system showed that OsS40-14 was 40.95% enriched in the transcription start site (TSS) of the genome. Sequence clustering analysis showed that OsS40-14 protein was mainly enriched and bound to TACCCACAAGACAC conserved elements. The seed region “ACCCA” of OsS40 proteins was identified by single nucleotide mutagenesis EMSA. The integrative analysis of transcriptome and CUT&Tag-seq datasets showed 153 OsS40-14-targeted DEGs, they mainly enriched in plastid organization and photosynthesis process at dark-induced condition in *oss40-14* relative to WT. Among them, eleven candidate targets of OsS40-14 such as Glucose 6-phosphate/phosphate translocator, Na+/H+ antiporter, Catalase, Chitinase 2, Phosphate transporter 19, OsWAK32, and OsRLCK319 were directly targeted and upregulated confirmed by ChIP-PCR and RT-qPCR. It demonstrates a novel model of OsS40-14 mediating macromolecule metabolism and nutrient recycling controls the plastid organization during dark-induced leaf senescence.

**Significant statement:** Involvement of OsS40-14 in macromolecule catabolism, nutrient recycling, and ROS homeostasis revealed a plastid organization defection of dark-induced senescence in rice

## Introduction

Leaf senescence is a natural developmental process at the final stage of leaf development; it involves nutrient and energy remobilization from resource to developing tissue, involving a series of changes in cellular physiological, biochemical, and molecular levels (Pyung *et al*., 2007; Guo, 2013). Leaf senescence can be triggered by abiotic stresses, such as darkness (Brouwer *et al*., 2012; Sobieszczuk-Nowicka *et al*., 2018; Gad *et al*., 2021). Numerous studies have shown that there is some overlap between naturally developmental senescence and dark-induced senescence. (Buchanan-Wollaston et al., 2005). Detailed analyses of gene expression patterns have revealed inconsistencies between the dark-induced and developmentally controlled processes (Becker and Apel, 1993; Buchanan-Wollaston *et al*., 2005; Van Der Graaff *et al*., 2006; Chen *et al*., 2010; Guo and Gan, 2012). Most of the genes that depend on hormones such as ethylene, abscisic acid (ABA), and jasmonic acid (JA) or ROS for expression during natural leaf senescence were also up-regulated during dark-induced senescence, indicating that these signaling pathways play an active role in both aging and dark-induced leaf senescence in Arabidopsis (Buchanan-Wollaston et al., 2005). In rice, among the 14 senescence-associated genes (SAGs) characterized, 11 genes were associated with both dark-induced and natural leaf senescence (Lee *et al*., 2001; Cao *et al*., 2022). Several age-dependent SAGs in rice have been demonstrated to regulate dark-induced leaf senescence, such as *NON-YELLOW COLORING1/3* (Os*NYC1/3*), *EARLY FLOWERING3.1* (Os*ELF3.1*), Os*NAP*, and DNA-binding with one finger 2.1 (Os*Dof2.1*), as well as MYB transcription factor (OsMYB102) (Kusaba et al., 2007a; Morita et al., 2009; Liang et al., 2014a; Sakuraba et al., 2016; Piao et al., 2019). The research of aging- and darkness-related leaf senescence genes has been greatly aided by high-throughput sequencing technologies.

Despite dark-induced leaf senescence (DILS) results in a clear loss of chlorophyll, disassembly of cellular elements and a lack of photosynthetic activity, none of which can be distinguished from the age-dependent natural senescence (Buchanan-Wollaston *et al*., 2002, 2005). Genes induced by dark-induced and age-dependent senescence still trigger different downstream nuclear gene expression profiles and signaling pathways (Breeze *et al*., 2011; Buchanan-Wollaston *et al*., 2005; Kanazawa *et al*., 2000). Previous research has shown that the photochromic-interacting factors (PIFs) signaling module is crucial in the dark-induced senescence of leaves (Liebsch and Keech, 2016; Gad et al., 2021). In Arabidopsis PIF4/PIF5 are essential transcriptional activators of dark-induced leaf senescence. Darkness induces them in a phytochrome B (phyB)-dependent manner. These two PIFs, along with ORE1, ETHYLENE INSENSITIVE 3 (EIN3), ABA INSENSITIVE 5 (ABI5), and ENHANCED EM LEVEL (EEL), finally regulate chloroplast maintenance, hormone signaling, chlorophyll metabolism, and senescence master regulators during dark-induced senescence *via* multiple coherent feed-forward loops (Sakuraba *et al*., 2014; Song *et al*., 2014; Qiu *et al*., 2015). Light deprivation can cause a decrease in photosynthesis and, as a result, carbon starvation. Some transcription factors involved in the low-energy response also play essential regulatory roles in the dark-induced senescence of leaves. For example, overexpression of BASIC LEUCINE ZIPPER63 (bZIP63) or bZIP1 that was responsive to low energy could accelerate dark-induced leaf senescence (Dietrich et al. 2011; Mair et al. 2015). During dark induced senescence conditions, autophagy participates in the movement of cellular components and serves as a quality assurance procedure by facilitating regulated degradation and recycling processes (Gad et al. 2021; Paluch-Lubawa et al. 2021). However, the detail mechanism of dark-induced rice leaf senescence still is limited understood.

The plant-specific senescence associated S40 family proteins contain a plant-specific domain of unknown function 584 (DUF584) are widely present in plants (Fischer-Kilbienski et al., 2010; Zheng et al., 2019). DUF548 is an intriguing C-terminal domain sharing the sequence GRXLKGR(D/E) (L/M)XXXR(D/N/T)X(I/V)XXXXG(F/I) which is highly conserved in plant species (Jehanzeb et al., 2017). AtS40s, OsS40s, and barley S40 (HvS40) participate in regulation of natural and stress-reduced leaf senescence (Krupinska et al., 2002; Fischer-Kilbienski et al., 2010; Jehanzeb et al., 2017; Zheng et al., 2019). Loss-of-function mutant of *AtS40-3* leads to a stay-green phenotype under both natural and dark-induced leaf senescence conditions (Fischer-Kilbienski et al., 2010). Moreover, AtS40.3 and HvS40 are assigned as DNA binding proteins (Krupinska et al., 2002; Fischer-Kilbienski et al., 2010). Our previous work has identified the S40 family including 16 members in rice, among them several members were involved in dark-induced leaf senescence, rice grain filling, and response to environmental cues (Habiba et al. 2021; Zheng *et al*., 2019). OsS40-14 is one of members of rice S40 family. It encodes a nuclear protein and is associated with natural and darkness-induced leaf senescence (Zheng et al., 2019; Habiba et al., 2021). OsS40-14 CRISPR/Cas9 editing lines (*oss40-14*) showed a stay green flag leaf and delayed senescence under dark-induced condition, and the *oss40-14* mutants have larger grains and high yield phenotype, indicating that loss of *OsS40-14* has beneficial traits for crop production in normal condition (Habiba et al. 2021). However, its molecular role in darkness-induced leaf senescence is unknown.

In this study, we further characterized OsS40-14 function in detached flag leaf under dark-induced condition and primary leaf of rice seedling under dark extensive condition using two independent *oss40-14* CRISPR/Cas9 editing lines and two *OsS40-14* overexpressing transgenic lines (*oe OsS40-14)* compared to wild type. Further, we developed a CUT&Tag (cleavage under targets and tagmentation) sequencing strategy to screen the direct targets of transcription factor in genome-wide level by using rice protoplast transient transformation assay. By using MEME program, the genome conserve binding site and of OsS40-14 was enriched, then the binding seed region was identified by electrophoresis mobility shift assay. The integrative analysis of the CUT&Tag dataset and the transcriptome of loss-of *OsS40-14* genotype relative to WT with darkness condition reveals that a novel model of the OsS40 family member is involved in macromolecule metabolism, nutrient recycling process, and affects plastid organization during dark-induced leaf senescence in rice.

## Material and methods

### Plant materials and growth conditions

The *oss40-14* mutant and its parental WT japonica rice variety Zhonghua 11 (ZH11) were grown in a growth chamber under 16 h light at 30°C/8 h night at 22°C in growth conditions. *OsS40-14* CRISPR transgenic plants *oss40-14* and overexpression lines (*oeOsS40-14*) were produced by the Biogle company (Hangzhou, China) using japonica rice variety Zhonghua 11. Genomic DNA from individual transgenic plants was isolated using Edwards buffer (Chen and Kuo, 1993) for PCR analysis. The PCR products were amplified with *OsS40-14* targeted-specific primers and were sequenced directly. The *OsS40-14* -specific primers were designed for amplifying targeted regions of *OsS40-14* (Supplementary Table S13). Sequences were decoded with DSDecode (http://skl.scau.edu.cn/dsdecode/) (Liu et al., 2015). The CRISPR-GE online tool (http://skl.scau.edu.cn/) was utilized to identify off-target sites for the target regions. Three putative off-target sites were found in intergenic regions, with no off-target sites identified in exon regions (Habiba et al., 2021).

### Darkness treatment

For the dark induced senescence (DIS) experiments, we used whole plants or detached leaf. 10-days-old WT and *oss40-14* mutants’ seedlings were incubated in complete darkness in Yoshida’s culture solution (Yoshida *et al*., 1971). The detached leaf segments from rice flag leaf grown under LD (14-h light/day) conditions were incubated on 3 mM MES (pH 5.8) buffer with the abaxial side up at 28 °C in complete darkness to induce DIS. The color changes of leaves were observed and photographed. The leaf segments were sampled at the specified days of dark incubation (DDI) for each experiment. Three biological replicates were used.

### Vector Construction

The full open reading frames of *OsS40-14* were amplified by PCR from cDNA pool. Appropriate restriction sites were included within the primers for subsequent cloning. Plasmids were obtained by using Novozyme’s "ClonExpress® II One Step Cloning Kit" (C112) kit to obtain the linearized vector and the target gene insert by homologous recombination with the adapter sequence in the infusion cloning manner. All clones were validated by sequencing. To generate *OsS40-14* overexpression line, *OsS40-14*cds was cloned into the KpnI-BamHI sites of a pUN1301 vector to obtain the pUN1301-OsS40-14 construct. OsS40-14 was driven by an ubiquitin promoter in the construct and a GUS marker was carried in the vector pUN1301 as described previously (Ge *et al*., 2004). For using in protoplast transformation, the expression vectors p2GWF7 (Karimi et al., 2002) with C-terminal GFP fusion of OsS40s-GFP were previously described by (Zheng *et al*., 2019). pGADT7 (AD) and pGBKT7 (BD) were used for the construction of yeast two-hybrid vectors. pRTVcVN (Accession No. MH373677) 0.1, referred to as VcVN; pRTVcVC (accession number MH373678.1) (He *et al*., 2018), VcVC; pRTVnVN (referred to as VnVN) and pRTVnVC (referred to as VnVC) for bimolecular fluorescence complementation experiments (BiFC). pCold vector (Hayashi and Kojima, 2008) was used for recombinant OsS40-14 protein expression in *E Coli*. Primers for all constructs generated in this study are listed in Supplementary table S13.

### Yeast two-hybrid assay

Experiments for Yeast Two-Hybrid (Y2H) assays were performed following the procedures outlined in the Yeast Protocols Handbook (Clontech). The respective combinations of GAD-fusion and GBD-fusion plasmids were co-transformed into yeast strain Y2H-Gold (Clontech) and colonies grown on synthetic defined (SD) medium with -Leu/-Trp and selected in SD medium with -Leu/-Trp/-Ade and higher stringency SD/-Leu/-Trp/-His/Ade plates. After 3 days of incubation at 28°C, the growth of each strain was measured. The transforms containing empty plasmids pGADT7 and pGBKT7 served as negative controls. Growth on synthetic medium-Trp-Leu-His-Ade indicated positive protein-protein interaction. Three biological replicates were performed for each combination in every growth assay.

### BiFC Assay

The VcVn- OsS40-14 and VcVc- OsS40-14 plasmids were extracted by the "EndoFree Plasmid Midi Kit" (CW2105), then were co-transformed into rice protoplasts (about 2.5 × 10^6^ cells/ml) prepared according to the description (Zheng *et al*., 2019) and incubated overnight at room temperature in darkness. Fluorescence signals were observed using a Leica TCS SP8 confocal laser scanning microscope. Leica TCS SP8 confocal software was used to process the images. Adobe Photoshop and Adobe Illustrator were used to organize the figures. Co-transfection of VnVn and VnVc empty vector was used as the negative control.

### Chlorophyll contents and Fv/Fm measurement

The chlorophyll content of the detached leaves was measured based on the method described previously (Lichtenthaler and Wellburn, 1983; Fatima *et al*., 2021). Chlorophyll was extracted from 2 leaf discs which mixed with 95% ethanol in a 1.5 milliliter (mL) Eppendorf tube. After incubating for 24 hours in the dark, pigments were extracted and the absorbance of extracted pigments was measured at 470, 649, and 665 nm using a spectrophotometer (L3, INESA, China), and the total chlorophyll concentration was calculated using the equations mentioned (Lichtenthaler and Wellburn, 1983). According to Shao’s description, the chlorophyll fluorescence was measured and the image was recorded using an Imaging-PAM-Maxi (Walz, Effeltrich, Germany) (Shao *et al*., 2008.). The minimum fluorescence at open PSII centers (Fo) and the maximal quantum yield of photosystem II (PS II; Fv/Fm) photochemistry was measured after adaptation to complete darkness for 20 min.

### Detection of Reactive Oxygen Species (ROS)

Hydrogen peroxide (H_2_O_2)_ and O^2−^ were detected with 3,3’-diaminobenzidine (DAB) and nitroblue tetrazolium (NBT) staining, respectively. H_2_O_2_ in the rice leaves was detected using DAB as previously described (Kim et al., 2018). Briefly, for DAB staining, rice leaves were dipped into a 1 mg/ml DAB solution (pH 5.7) and incubated at room temperature in the dark for around 8 hours. After that, the leaves were distained three times with 95% ethanol and heat in a water bath at 95°C for 15 minutes. For NBT staining, rice leaves were infiltrated with 0.5 mg/ml NBT solution (pH 7.5) after which the leaves were maintained overnight at room temperature. The leaves were submerged in absolute ethanol, heated in a boiling water bath for 10 minutes, cooled for 30 minutes at room temperature, transferred to paper, and photographed three times against a contrast background to remove the staining. An Epson Perfection V600 Photo scanner (Epson China, Beijing, China) was used for the imaging process.

### Transcriptome analysis

Total RNA for the RNA-seq analysis was extracted from leaves of WT and *oss40-14* plants incubated in complete darkness for 3 days. One biological replicate from *oss40-14.1* independent mutant and other two biological replicates from *oss40-14.2* independent mutant were performed for RNA seq analysis as both mutants exhibit a dark-induced phenotype. RNA-Seq was performed with an Illumina HiSeqX instrument (Illumina. San Diego, CA, USA). Using HISAT2 software (version 2.1.0), the trimmed reads were mapped onto the reference rice genome MSU7 (RGAP, http://rice.plantbiology.msu.edu/, build 7.0) after the low-quality bases (Q 20) and short sequence reads (length 20) were removed. Raw read counts were estimated using feature Counts software and normalized with R package, DESeq2 (Love *et al*., 2014) (R version: 3.5.0, DESeq2 version: 1.22.2). The DEGs were estimated with the software package DESeq2, and the genes that exhibited p-value < 0.05 and absolute log2 (fold change) ≥1 were determined to be significantly differentially expressed. Gene ontology (GO) and KEGG enrichment analyses were performed using the graphical enrichment tool Shiny Go v0.61(Shannon *et al*., 2003). The function categories of the genes were selected with the enrichment FDR (false discover rate) of p < 0.05.

### CUT&Tag assay

CUT&Tag experiment was performed in rice protoplasts. The expression vector *p2GWF7-OsS40-14-GFP* or empty vector *p2FGW7-GFP* was transfected into protoplasts isolated from the green leaf sheaths of 2-week-old rice seedlings as described previously (Zheng et al. 2019). After 12h incubation, 3 ml transformed protoplasts were collected and treated with 0.1% formaldehyde for 2 min. Followed by cross-linking, the harvested protoplasts were resuspended in 1 ml Nuclei isolation buffer (100mM MOPS pH 7.6, 10mM MgCl2, 0.25M sucrose, 3% Dextran T-70, 2.5% Ficoll 400, 1mM DTT). After centrifugation, the collected nuclei of protoplasts were used for CUT&Tag experiments according to the instructions of the commercial kit (HyperactiveTM In-Situ ChIP Library Prep Kit for Illumina, TD901-TD902, Vazyme, China). The mouse anti-GFP antibody (Sigma, G6795) was used as the primary antibody (1:100) to incubate with ConA beads-treated nuclei for at least 4 h or at 4 °C overnight. After the CUT&Tag reaction, the DNA fragments were isolated for PCR enrichment and Next-Generation Sequencing (NGS). Reads containing adapters and low-quality reads are removed through quality control to obtain clean data. Clean data was analyzed using the "CUT & Tag_ tool" tool in the Novozymes bioinformatics cloud platform (http://cloud.vazyme.com:83), GFP library served as a control. The bw file was generated using the "bam2bigWig" tool in Novozymes bioinformatics cloud platform, visualized and analyzed by IGV software. The sequence of binding motif was clustered using MEME suite program (https://meme-suite.org).

### Quantitative Real-Time PCR

Total RNA was extracted from leaves using TRIzol Extraction reagent (Invitrogen), according to the manufacturer’s protocols. The first-strand cDNA was obtained from 1 µg of total RNA, using the Synthesis Kit (Thermo Fisher Scientific), eliminating the contaminant genomic DNA. qRT-PCR was performed to analyze the expression of genes in the CFX96 machine (Bio-Rad Company, Hercules, CA, USA) in a whole volume of 15 µl, using SYBR Green Master (ROX) (Newbio Industry, China) as per the instructions of the manufacturer. The endogenous *OsACTIN* gene (LOC_Os03g50885) and *OsUBQ5* (LOC_Os01g22490) were used as internal controls. Relative expression levels of target genes were calculated using the 2−ΔΔCt method for multiple reference gene as mentioned (Riedel *et al*., 2014). In this experiment three biological replicates were tested, and each biological replicate contains leaves from three independent plants. Primers used for this experiment are listed in Supplementary Table S13.

### ChIP-qPCR

For the chromatin immunoprecipitation (ChIP) assay, the *35S:OsS40-14-GFP* and *35S:GFP* plasmids were transfected into rice protoplasts as previously described (Habiba *et al*., 2021). The protoplasts were then subjected to crosslinking for 20 min with 1% formaldehyde under vacuum. The chromatin complexes were isolated and sonicated as previously described (Saleh *et al*., 2008). An anti-GFP polyclonal antibody (Abcam) and Protein A agarose/salmon sperm DNA (Millipore) were used for immunoprecipitation. After reversing the crosslinking and protein digestion, the DNA was purified using a QIAquick PCR Purification kit (Qiagen). Enrichments of the selected promoter regions of candidate genes were resolved by comparing the amounts in the precipitated (IP) and nonprecipitated (INPUT) DNA samples, which were quantified by qPCR using designed region-specific primers (Figure 9G; Supplemental Table S13).

#### Recombinant OsS40-14 protein preparation

pCold-OsS40-14-GST and empty pCold vector plasmids were transformed to Rosetta (DE3) strains, at 16^0^C, 0.5mM IPTG induction for 16 hrs, then extracted and purified the OsS40-14 protein with GST beads. The purified OsS40-14 recombinant protein was detected by western blot using an antibody against GST (Abcam) and was stored at -80^0^C and used for EMSA assay.

#### Electrophoresis mobility shift assay (EMSA)

The DNA probe preparation: the linker DNA fragment with Cy5 labelled 5’-GGTCTGGTCCCTGTG-Cy5-3’ and the DNA probe fragments or series mutagenized probes of F chain CCAGACCACGGGCAC and R chain were artificial synthesized by Shangya Bio company. The three DNA fragments were incubated with the ratio of volume (3:3:1) at 95^0^C for 5 min, then turned to room temperature for 2-3 hrs.

According to the method described by Miao (Miao et al., 2013). Briefly, mix the Cy5 labeled DNA probe, the purified OsS40-14 recombinant protein and 5x binding buffer incubated on the ice for 2hrs, then the samples were loaded and electrophoresed on 6% acrylamide PAGE-gel, non-labeled DNA probe was used a competitor. After running, the gel was detected with BOX Chemi XT4 machine scanning and photographing.

### Statistical Analysis

The data in all figures were determined by at least three biological replicates. To determine statistical significance, one-way ANOVA followed by Tukey’s HSD test or student’s t-test was performed. Different letters represent statistical significance, and P<0.05 indicates a significant difference. All these were carried out using the GraphPad Prism software version 8 (GraphPad Software, San Diego, CA, USA).

## Results

### OsS40-14 is highly expressed in phloem cells and enhanced by dark induction

*OsS40-14* highly expressed in leaf and root (Habiba et al., 2021). To insight which kind of cell type OsS40-14 existed, a screening of public single-cell transcriptome databases of rice leaf and root (Zhang et al., 2021; Wang et al. 2022) was performed. The expression of *OsS40-14* was highly clustered in the mestome sheath cells of leaf (Figure 1A) under stress condition. Quantitative real-time PCR analysis of rice leaves demonstrated a significant induction of *OsS40-14* transcript levels in both detached primary leaves and flag leaves after two days of darkness treatment (Figure 1B), suggesting that OsS40-14 may function as a dark-induced senescence-associated factor.

**Figure 1.**
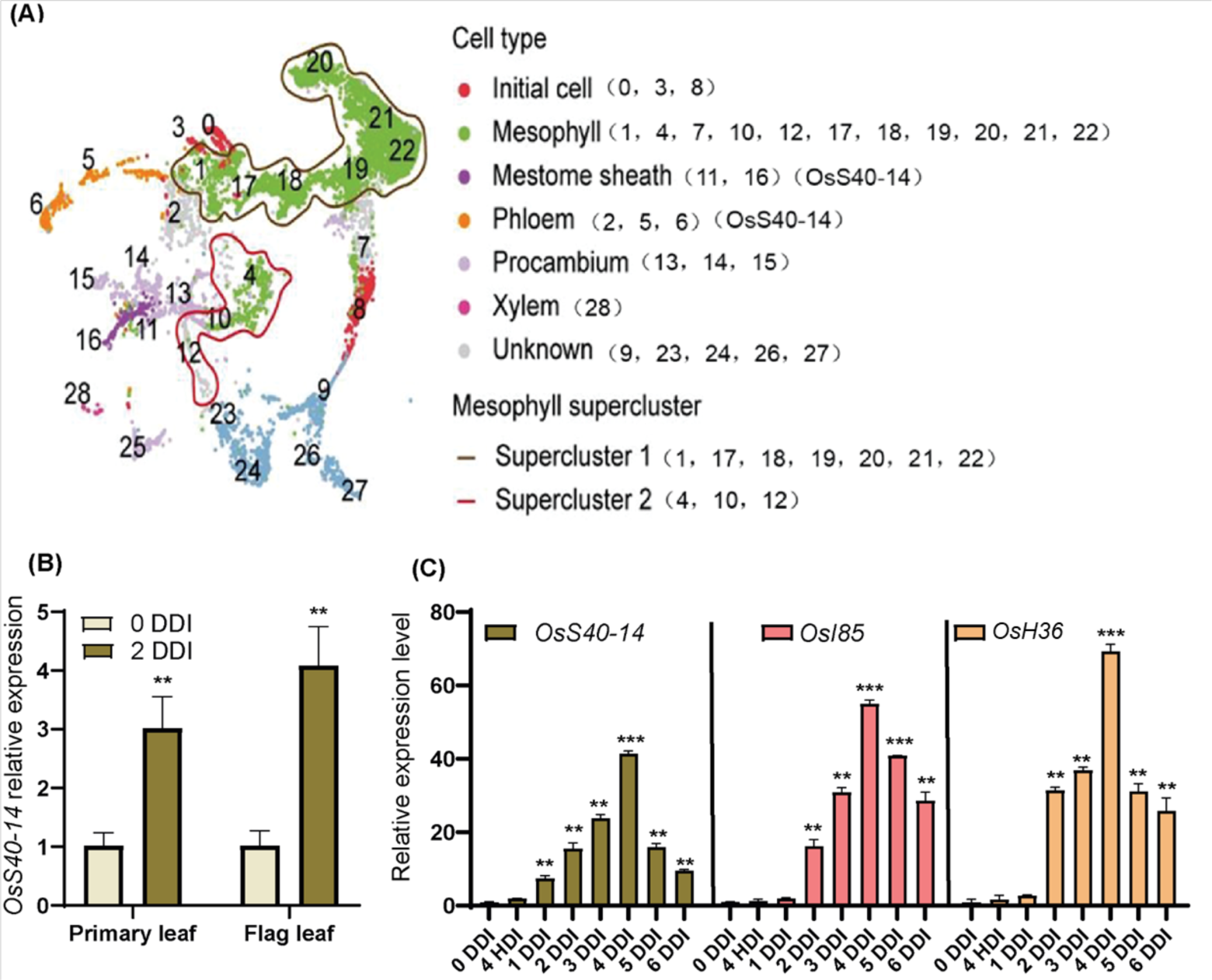
Expression profiles of *OsS40-14* and senescence-associated genes in rice during dark-induce senescence (DIS) condition compared to natural condition. (A) The expression of *OsS40-14* was highly clustered in the mestome sheath cells of leaf and root by screening of public single cell transcriptome databases of rice leaf (Wang *et al*., 2021; Zhang *et al*., 2021; Liu et al., 2021; Zong et al., 2022). The number indicates the cell type of rice leaf and root referred to Wang et al., (2021); (B) The relative transcript levels of *OsS40-14* in detached primary and flag leaf discs from the japonica cultivar” Zhonghua 11”; (C) Expression of *OsS40-14, OsI85, and OsH36* measured in 10 days old rice seedling after complete darkness treatment for 0DDI to 6 DDI at 28 °C. *OsS40-14* mRNA levels were determined by RT-qPCR analysis and normalized to that of *OsACTIN* and UBQ. Mean and SD values were obtained from at least three biological samples. Different letters indicate significant differences among DDI according to one-way ANOVA and Tukey’s multiple comparison test (p<0.01).(B, C) DDI, day(s)of dark incubation.

To further substantiate these findings, a detailed time-course expression experiment was performed on 10-day-old rice seedlings. Remarkably, a night extension of up to 24 h leads to a seven-fold accumulation of *OsS40-14* transcripts (Figure 1C), which further increased during an extended night by up to 41-fold until 5 days of dark incubation (DDI). Other three members of OsS40 family *OsS40–1*, *OsS40–2*, *OsS40–12* were also upregulated in the seedlings at late stage of extended night after dark treatment (Supplementary Figure S1). Senescence-associated genes (SAGs) related to dark-induced senescence (DIS) in rice, including two dark induced marker genes, glyoxylate aminotransferase (*OsH36*) and isocitrate lyase (*Osl85*) (Lee et al., 2001) were similarly upregulated with prolonged dark incubation (Figure 1C). These results indicate that OsS40-14 play a role in the dark-induced response of rice.

### OsS40-14 can form homomeric oligomer

OsS40-14 protein shares a conserve DUF584 domain with OsS40 family. It was located in the nucleus and has transcriptional activation activity (Zheng et al., 2019). In order to study their molecular function, we detected their interaction by yeast two-hybrid assay firstly. OsS40-14 and OsS40-14 were fused with the Gal4 DNA-binding domain (in pBD bait vector) and activation domain (in pAD prey vector), Upon co-transformation of *AD-OsS40-14* with *BD-OsS40-14* into AH109 yeast cells, the growth on selection media deficient in Leu, Trp, His, and Ade indicated the detection of homomeric oligomers of OsS40-14 (Figure 2A). When the empty prey or bait vectors were transformed with AD-OsS40-14 or BD- OsS40-14, respectively, the colonies did not grow in the selected medium, whereas the positive control grew well, suggesting that OsS40-14 can form homooligmers in yeast cells.

**Figure 2.**
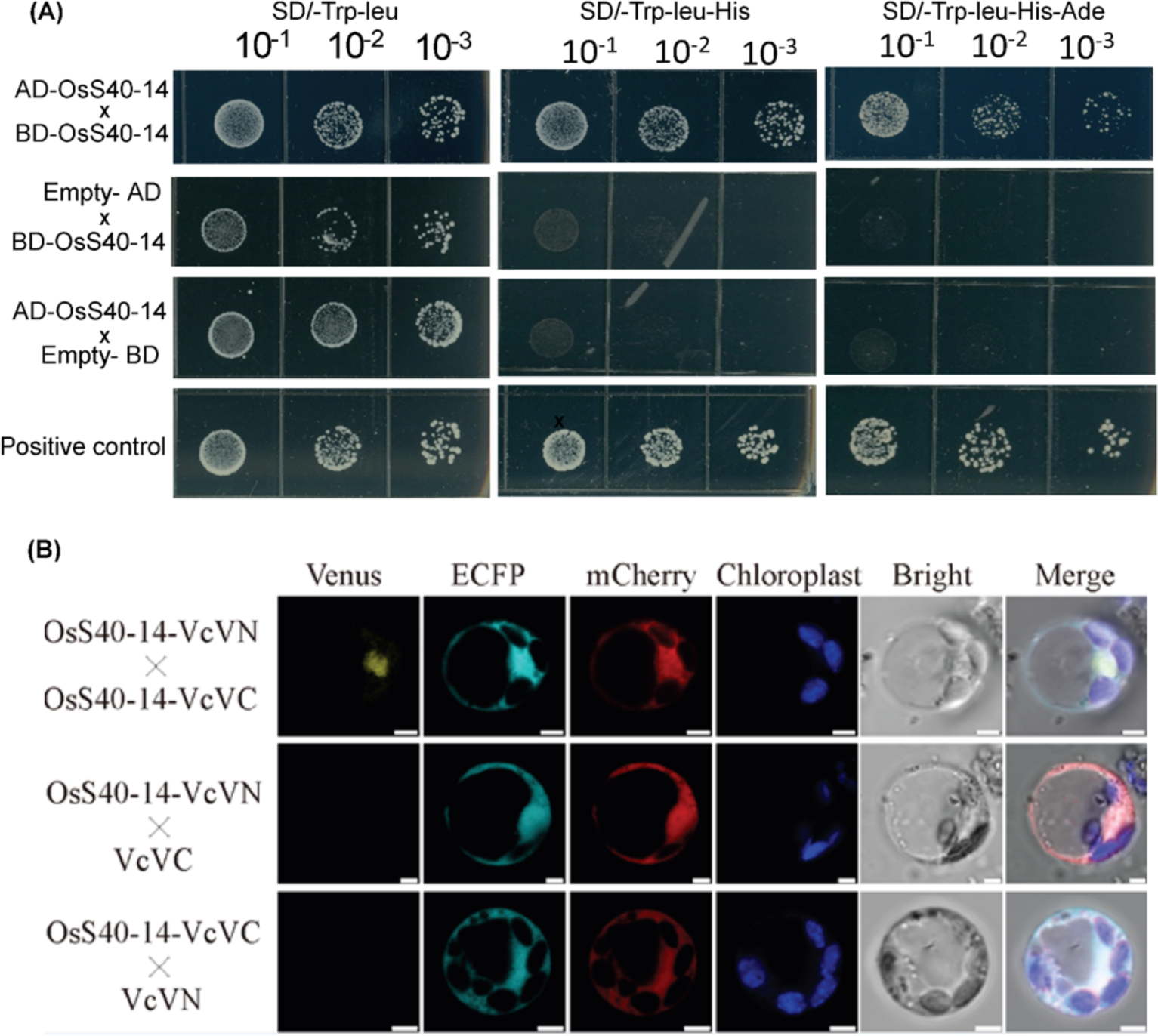
Homomeric interaction detection of OsS40-14 by yeast two-hybrid and bimolecular fluorescence complementation (BiFC) assays.(A) Confirmation of OsS40-14 homomeric interaction in a yeast two-hybrid assay. Transformed yeast cells were cultured on synthetic defined (SD)/ -Trp-Leu, (SD)/Trp-Leu-His, (SD)/-Leu-Trp-His-Ade medium. The AD or BD empty vector was used as a negative control, and the interacting pair UPL5-WRKY53 was used as a positive control (B) The homomeric interaction detection of OsS40-14 in rice protoplasts by BiFC assay. Empty vector is used as Negative control.

To validate the interaction between OsS40-14 and OsS40-14 in planta, OsS40-14 was fused with N- or C-terminal parts of a split yellow fluorescent protein (YFP) for bimolecular fluorescence complementation (BiFC) assays. The plasmids of OsS40-14-VcVn and OsS40-14-VcVC were co-transformed in rice protoplast via PEG- mediated transformation. The detection of Venus fluorescence indicated the interaction of OsS40-14-VcVn with OsS40-14-VcVC in the nuclei of rice protoplast (Figure. 2B). As the negative controls, co-expression of OsS40-14-VcVn and the Empty-VcVc vector as well as Empty-VcVn and OsS40-14-VcVc did not enable the cells to produce the Venus fluorescence signal. (Figure 2B). These results indicate that OsS40-14 can physically interact with OsS40-14 forming homomeric oligomer.

### *OsS40-14* Crispr/Cas9 editing lines exhibited a stay green phenotype under darkness condition

In the previous works, the CRISPR/Cas9 editing lines of *oss40-14.1* were constructed and identified (Habiba *et al*., 2021). The *oss40-14.1* editing lines exhibited a slight stay green flag leaf and high grain weight agronomy traits (Habiba *et al*., 2021). In this study, we generated another independent CRISPR/Cas9 editing mutant line *oss40-14.2* to verify the phenotypic effects observed in the *oss40-14.1* mutant line. The *oss40-14.1* was the homozygous mutant with one base pair ‘G’ deletion in the target site of the exon, while *oss40-14.2* was the homozygous mutant with 2 base pairs ‘GA’ deletion in the target site of the exon (Supplementary Figure S2A, B). Incubation of the two mutants in darkness resulted in a delayed senescence of the detached flag leaf pieces and the whole *oss40-14* seedlings, although *OsS40-14* genes CRISPR/Cas9 editing genotype did not show significantly stay green primary leaf senescence phenotype in natural growth condition. The detached leaves of *oss40-14* and WT were treated for 5-day dark induction (DDI), the detached leaves of *oss40-14* were stayed green up to 3DDI compared to WT becoming yellow after 2DDI (Figure. 3A). The chlorophyll content was maintained high amount in detached leaves of the *oss40-14* at 72 hours’ dark treatment while it dropped significantly after 48 hours in WT (Figure. 3B). The distribution of chlorophyll fluorescence in the dark-induced condition was analyzed to observe photosynthetic capacity using an Image-PAM (Pulse-Amplitude Modulation) measuring system, as shown in Figure 3C. The maximum photochemical efficiency of photosystem II (Fv/Fm) significantly decreased in WT compare with *oss40-14* after 3DDI (Figure. 3C-D).

**Figure 3.**
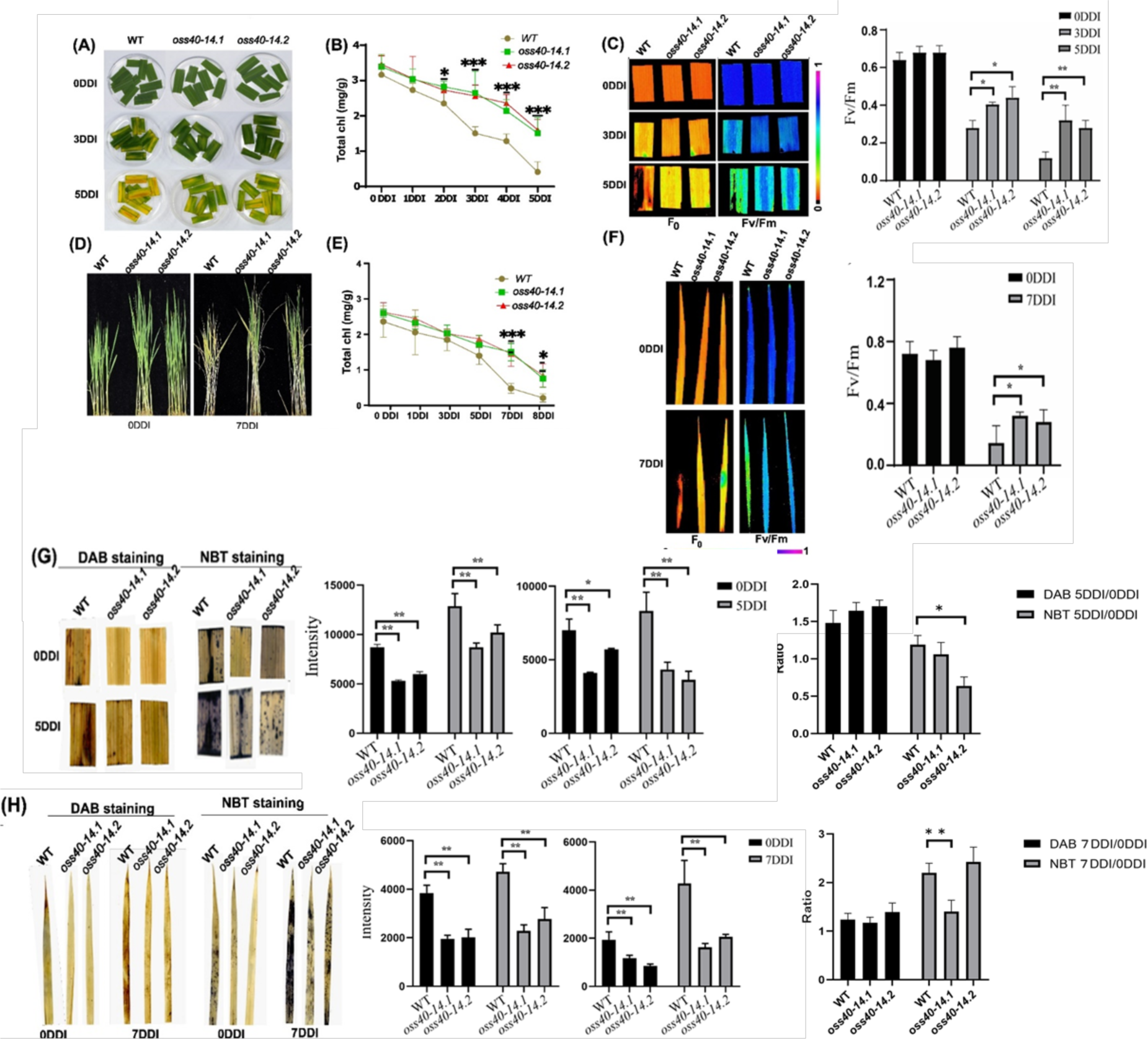
The *oss40-14* mutant exhibited stay green phenotype during DIS (A D) Phenotypes of dark-induced leaves from flag leaf segments and 10 days old seedling of WT and *oss40-14*. Leaf segment were incubated on 3 mM MES (pH 5.8) buffer with the abaxial side up at 28 °C in darkness.10-days-old seedlings were grown LD (14-h light/day) conditions and then were transferred to darkness at 28 °C for 7 days (7 DDI) (B, E) The changes of total Chl level (C F) The fluorescence images of the whole plants and leaf segment of WT and *oss40-14* mutants. The fluorescence images are taken by Image-PAM using the plants after 30 minutes dark-adapted (G, H) H_2_O_2_ levels and O^2-^ levels detected by DAB and NBT staining with photos and digital calculation and ratio of Image J in dark-induced flag leaf segments and 10-days-old seedlings of *oss40-14* mutant and WT. Asterisks indicate significant difference compared with the expression level of *OsS40-14* at 0 DDI (B, E) (Student’s t-test, *P<0.05, ***P<0.01) (A, D) DDI, day(s) of dark incubation.

Next, this stay green phenotype of *oss40-14* mutant was also examined using 10 days old whole seedlings. After 7 DDI, many leaves of WT seedlings became yellow, while most *oss40-14* leaves retained their green color (Figure 3E). The leaf segments of *oss40-14* plant retained their green color longer than the WT leaves, showing a striking difference at 7 DDI (Figure 3E). Consistent with the visible phenotype, *oss0-14* mutants had higher chlorophyll (Chl) levels and higher photosynthetic capacity (Fv/Fm), similar to detached leaf segments (Figure 3F, G, H).

We further detected the senescence associated parameter ROS status of detached flag leaves and primary leaves after dark treatment to investigate the retained green phenotype in *oss40-14* and WT plants. We stained for hydrogen peroxide (H_2_O_2_) and superoxide radical (O^2−^) using DAB and NBT staining assays, respectively. The signal status of DAB and NBT staining was weakened in *oss40-14* leaves compared to ZH11 WT (Figure 3I, J). The digital counts of H_2_O_2_ and super oxygen and their ratio of dark treatment relative to non-treatment revealed that the contents of H_2_O_2_ and super oxygen were significantly decreased in the *oss40-14* leaves and the ratio of super oxygen content was pronouncedly increased in WT than in the *oss40-14* leaves after darkness treatment, but the ratio of H_2_O_2_ did not significantly altered after darkness treatment (Figure 3K), suggesting that OsS40-14 was involved in balance of super oxygen of ROS and tolerance to darkness under dark-induced leaf senescence condition.

Further to genetically confirm this issue that the overexpression of *OsS40-14* induces early leaf senescence, we generated transgenic rice lines overexpressing *OsS40-14*. RT-qPCR analysis revealed that OsS40-14 was significantly upregulated in both overexpression line (*OE-3-1*, *OE-2-3*) transgenic lines (Supplementary Figure S3A). In contrast to the mutant phenotype, the leaf segments of two *OsS40-14* overexpression lines exhibited an early yellowing phenotype at 4 days of dark incubation compared to the wild type (Supplementary Figure S3B-C). These findings collectively support the notion that OsS40-14 plays a role in accelerating dark-induced leaf senescence in rice.

### Loss of *OsS40-14* altered nuclear gene reprograming of metabolic, photosynthetic, rhythmic, and response to stresses under dark-induced condition

To understand the molecular mechanisms of OsS40-14-regulated leaf senescence under dark-induced conditions, we constructed genome-wide gene expression profile in the detached leaves of *oss40-14* and WT plants after 0DDI and 3DDI using RNA sequencing (RNA-seq). Total three biological replicates of two independent lines were prepared. Evaluation of the two replicates of RNA-seq datasets were analyzed. Four treatments were processed twice, resulting in a total of 8 transcriptome expression datasets. The transcriptome expression data were reduced to two dimensions using principal component analysis (PCA) method. The first principal component (PC1) showed that, except for slight separation between the two replicates of the wild-type treatment, the replicates of the other treatments had good reproducibility, especially the two dark-treated materials; the data separation between different treatments was obvious, with good discriminability (Figure 4A). Based on the criteria of absolute log2(fold-change) value ≥1 and adjusted p value <0.05, a total of 1557 differential expressed genes (DEGs) (Supplementary Table S1), including 955 upregulated genes and 602 downregulated genes, were identified in the *oss40-14* compared to those in WT under normal condition (Figure 4B, Supplementary Table S2). while 63 DEGs, including 36 upregulated genes and 27 downregulated genes, were identified in the *oss40-14* compared to those in WT after darkness treatment (3DDI) (Figure 4B; Supplementary Table S3). When the comparative analysis of 3DDI-DEGs relative to 0DDI-DEGs in the *oss40-14* compared to WT were performed, based on the criteria of a greater than log2≥1-fold change with significance at false discovery rate adjusted p < 0.05, a total of 1585 dark-induced differentially expressed genes (Di-DEGs) were identified in *oss40-14* relative to WT(ZH11) under dark treatment(3DDI) compared with control conditions (0DDI) (Figure 4C; Supplementary Table S4-5). Gene ontology (GO) analysis of 1585 Di-DEGs enrichment in biological processes showed that genes related to photosynthesis, plastid organization, small molecule metabolic process, and in response to stress stimulus process, as well as circadian rhythmic process were most significantly enriched in the *oss40-14* relative to WT (Figure 4D). It suggests that the function of *OsS40-14* gene is closely related to rice’s response to abiotic stress signals and the connection between photosynthesis and chloroplast metabolic processes. Among the genes associated with the function of *OsS40-14*, there is a significant enrichment of known stress-responsive transcription factor downstream target genes, including AP2/ERF/ABI3, NAC, bHLH, bZIP, and WRKY factors (Figure 4E).

**Figure 4.**
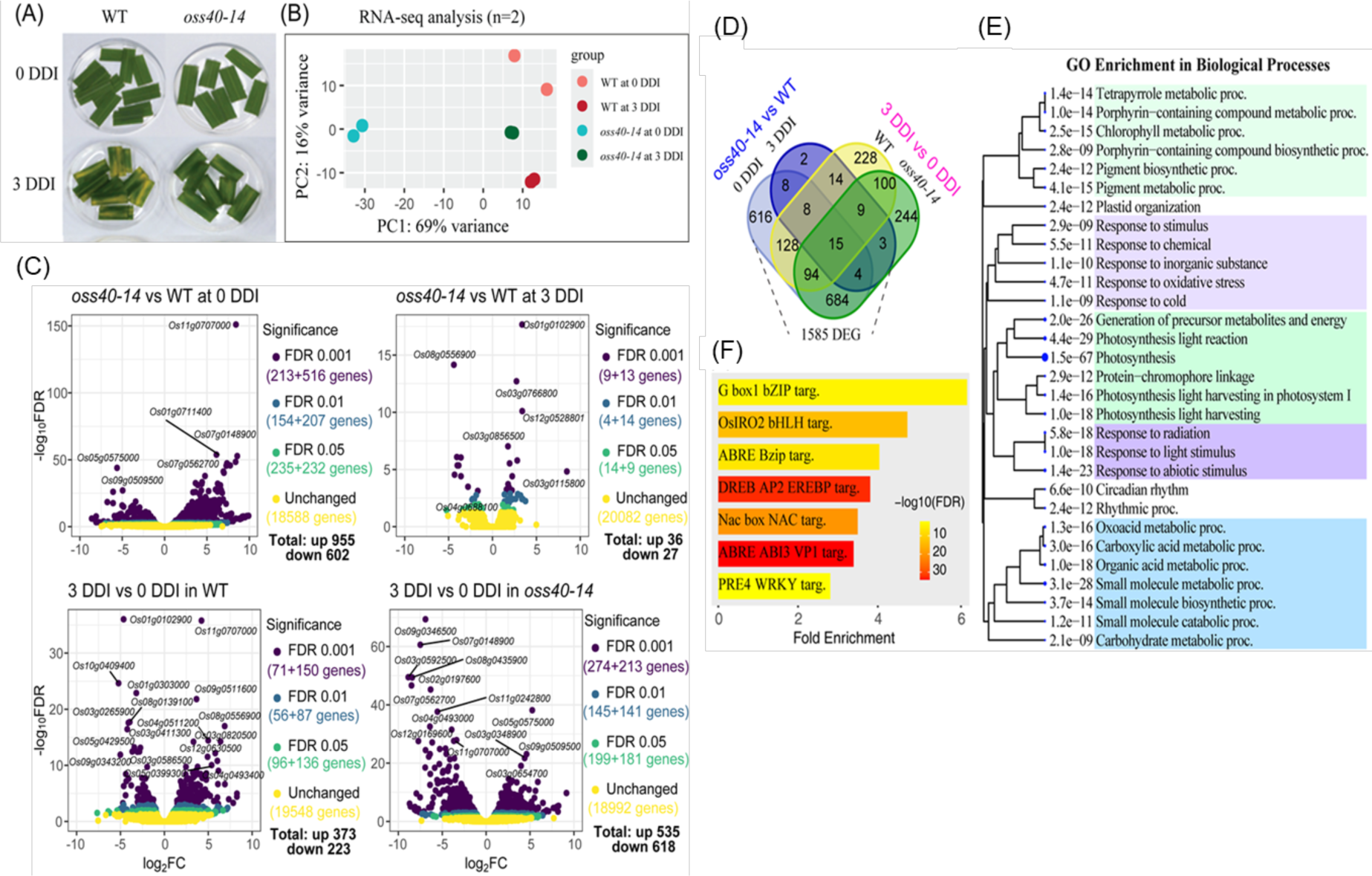
Darkness-induced yellowing senescence and transcriptome detection in the detached rice leaves.(A) Changes of rice detached leaves in dark treatment; wild type and mutant were photographed and sampled before treatment (0 DDI) and after 3 days of darkness treatment, respectively, and the experiment was repeated three times, with photographs of group 3 materials; (B) Sample variability of the results of the RNA-seq analysis and the correlation between two replicates. (C) Volcano plots of four transcriptomes analyzed for differentially expressed genes (DEG). The upper panels show the DEG results of mutant *oss40-14* relative to the wild type (ZH11) before (0 DDI) and after 3 days of treatment (3 DDI), and the lower panels show the DEG results of the wild type and *oss40-14* mutant after 3 days of dark treatment (3 DDI), respectively, relative to before treatment (0 DDI). (D) Venn diagram of the four groups of differentially expressed genes showing the distribution of DEG; (E) The head 30 GO: Biological Processes entries enriched by DEG 1585, with similar entries clustered together, the size of the dots indicates the significance of the enrichment, and the value is the FDR of its enrichment; (F) The database of known rice stress response transcription factors (STIFDB V2.0) downstream target genes enriched in DEG 1585; enrichment analyses of (E) and (F) in the figure were performed using Shiny GO 0.80 (run online on 2024-05-30).

### Loss of *OsS40-14* downregulated the gene expression related to several transcription factors and hormones under dark-induced condition

The above analysis reveals that the associated genes of transcription factor OsS40-14 are largely regulated by several classes of stress response transcription factors. To screen these associated genes for transcription factors and to explore the potential nodes in the downstream gene network of OsS40-14, the known gene set of the rice whole-genome transcription factor database (PlantRegMap/PlantTFDB v5.0, planttfdb.gao-lab.org) was downloaded and overlapping comparisons were used to obtain 74 transcription factors (Figure 5A; Supplementary Table S6).

**Figure 5.**
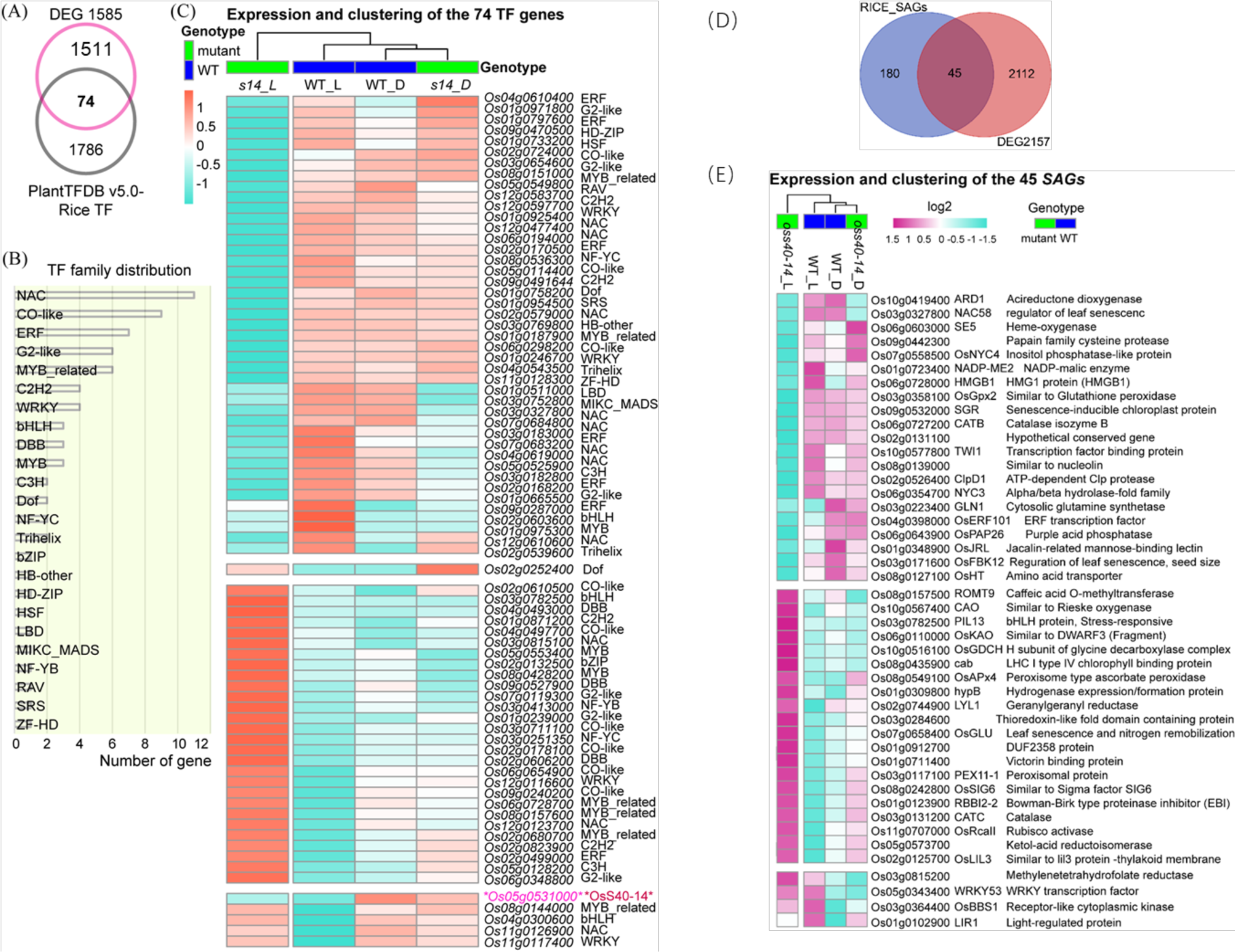
Transcription factor (TF) families and senescence-associated genes identified in the DEG dataset of *oss40-14* relative to wild-type. (A) Venn diagram showing the presence of 74 TF genes in 1585 DEGs, Plant Transcription Factor Database Rice Transcription Factor Set (PlantTFDBv5.0-rice TF downloaded on 2024-05-31); (B) The 74 transcription factors of rice are distributed in multiple plant transcription factor families; (C) The expression of 74 transcription factors of rice in four conditions (samples), OsS40-14 was used as the expression change comparison, but as a result of CRISP gene editing, the expression in the mutant only represents the activity of its promoter, and the heatmap was produced using programs such as R package heatmap; (D) Venn diagram showing the presence of 45 SAGs in 1585 DEGs. Plant senescence associated genes database 248 Rice SAGs (https://ngdc.cncb.ac.cn/lsd/); (E) The expression of 45 SAGs of rice in four conditions (samples), the heatmap was produced using programs such as R package heatmap.

Among these transcription factors, the top seven classes with the highest number of genes were NAC, CO-like, ERF, G-like, MYB-related, C2H2, and WRKY family members, all containing four or more genes (Figure 5B). Gene expression clustering analysis (k = 4) revealed that the up-regulated and down-regulated proportions of these transcription factors in the *oss40-14* mutant were approximately equal, at 44% and 56%, respectively (Figure 5C). For comparison, the expression data of *OsS40-14* gene was also added to the expression heatmap, but it should be noted that the *oss40-14* mutant is a CRISPR/Cas9 editing construct (base deletion and insertion) (Habiba et al., 2021), and the expression levels in the two sample materials of the mutant represent the activity of its promoter and cannot be considered as the expression level of the normal function protein gene. Through expression clustering, it was found that four transcription factors had expression changes consistent with *OsS40-14* gene expression in the wild type but opposite in the mutant. Additionally, there is an opposite trend in expression changes among members of the same gene family (Figure 5C), suggesting the following possibilities: (1) OsS40-14 may have specific regulation of different members; (2) OsS40-14 may interact with different nuclear factors at different gene loci, affecting the expression level of the corresponding genes; (3) the expression of different members of the same gene family is affected by OsS40-14 to different degrees, and the effects of other unknown factors are greater.

Phytohormones, such as abscission acid (ABA), ethylene, jasmonate acid (JA), salicylic acid (SA), strigolactone, gibberellic acid (GA), Cytokinin (CK), and brassinosteroids promote or inhibit plant senescence (Kusaba *et al*., 2013; Guo et al., 2021). It is noteworthy that 25 DEGs related to phytohormone biosynthesis and signaling categorized into ABA, JA, SA and ethylene were significantly downregulated in *oss40-14* mutant compared to wild type (Supplementary Table S7). After dark treatment, 18 phytohormone-related DEGs have been enhanced down-regulation in the *oss40-14* mutant compare to WT during dark induced senescence condition at 3DDI which includes ABA, Jasmonic acid, ethylene and auxin signaling (Supplementary table S7). For example, among them, the downregulated DEGs include *OsNAP,* encoding a senescence-associated NAC TF (Liang *et al*., 2014), *OsCCD1* encoding chlorophyll catabolic enzyme (Park *et al*., 2007), *OsLOX2* (Huang *et al*., 2014), and *OsLOX8* encoding (lipoxygenase) upregulate JA levels (Shim *et al*., 2019.) were down-regulated in *oss40-14* plants, even in non-senescent leaves (Supplementary Table S7). Different stress/ABA-activated protein kinases like OsSAPK2 (Lou *et al*., 2017), OsSAPK9, OsSAPK6 (Yu *et al*., 2022), a kind of SNF1-related protein kinase 2 (SnRK2), which are involved in the ABA signaling and abiotic stress tolerance, also significantly downregulated in *oss40-14* mutant compare to WT. Gene related to ABA signaling OsPYL1 related to dark induced senescence (Lee et al., 2015) was 6 times downregulated. *OsSLC1* (Lv *et al*., 2020) and *OsPrl5* (Spielmeyer *et al*., 2002), two gibberellin (GA)-related genes mainly related to the developmental process, are downregulated in the *oss40-14* mutant, however the cytokinin-related gene *OsLOG1*(Chen *et al*., 2022) is upregulated in the *oss40-14* mutant compare to WT. This result reveals that OsS40-14 regulates or interacts TFs and phytohormone related genes to control dark induced leaf senescence in rice.

### Loss- of *OsS40-14* affected 45 senescence associated genes (SAGs) up/down expression under dark-induced condition

Furthermore, 1585 differentially expressed genes were overlapped with the 248 known aging genes in rice (https://ngdc.cncb.ac.cn/lsd/) (Cao *et al*., 2022)), and 45 rice aging genes were associated with OsS40-14 (Figure 5D). Among them, 21 SAGs were found to be upregulated, and 24 SAGs were downregulated (Figure 5E; Supplementary Table S8). Of these, 21 SAGs, including genes related to phytochrome-interacting factor (*OsPIL13*), ASCORBATE PEROXIDASE 4 (*OsAPX4*), Catalase (*OsCAT*), *LTS1*, LEAF TIP SENESCENCE 1(*OsNaPRT1*), LESION AND LAMINA BENDING (*LLB*), ABNORMAL CYTOKININ RESPONSE 1 (*Fd-GOGAT1*), glycine decarboxylase H (*OsGDCH*), Light-harvesting like protein (*OsLIL3*) were significantly up-regulated in *oss40-14* mutant and interestingly these genes are involved in the senescence delaying process. In contrast, among the 24 SAGs, genes implicated in chlorophyll degradation, such as *OsSGR*, senescence-activated genes, such as NAC PROTEIN (*OsNAP*), *OsWRKY93* (Li et al., 2021), NAC domain-containing protein 11 (*OsNAC11*), *OsFBK12* were significantly downregulated in *oss40-14* mutant. Furthermore, nutrient recycling-related genes, including NADP-MALIC ENZYME 2 (*OscytME1*), EARLY RESPONSIVE TO DEHYDRATION1 (*OsClpD1*), expressed protein (Os09g0363500), peptidase A1 domain containing protein (Os02g0730700), and ORYZAIN GAMMA CHAIN (*Oryzain γ*) were also included in the downregulated SAGs in *oss40-14* compared to wild type. Our findings suggest that there is a significant regulatory role of OsS40-14 with upregulating the series of SAGs related chlorophyll degradation, macromolecule metabolic process, and nutrient recycling, with downregulating the SAGs related photochromic and ROS metabolic process during dark-induced leaf senescence.

### CUT&Tag-seq analysis reveals OsS40-14 targeting the conserve element with “ACCCA” seed region

Previous studies have shown that OsS40-14 localized in the nucleus and has transcriptional activity (Habiba *et al*., 2021). Given it is a transcription factor, OsS40-14 might directly bind to downstream target genes and affect their expression. To investigate the genome-wide profiling of OsS40-14 binding sites in rice cells, CUT&Tag assay combined with rice protoplast-based transient expression system was developed and performed for overexpressing OsS40-14-GFP and GFP control to identify its direct target genes (Supplementary Figure S5A; Supplementary Table S9). A total of 2311 putative targets were collected in OsS40-14-GFP relative to GFP. The distribution of CUT&Tag density in the region of the gene body (only 0.41%) and its upstream 5-kb flanking region (40.95%) revealed that the signals from CUT&Tag OsS40-14-GFP had higher intensity near the transcription start site (TSS) and in the gene body than those from CUT&Tag GFP control (Figure 6A-C; Supplementary Figure S5B-C). By using MEME program, the sequences of OsS40-14 targeting are clustered to three conserve elements: TACCCACAAGACAC, CGGTTATGG, and TATTCGAATAGCCG (Figure 6D).

**Figure 6.**
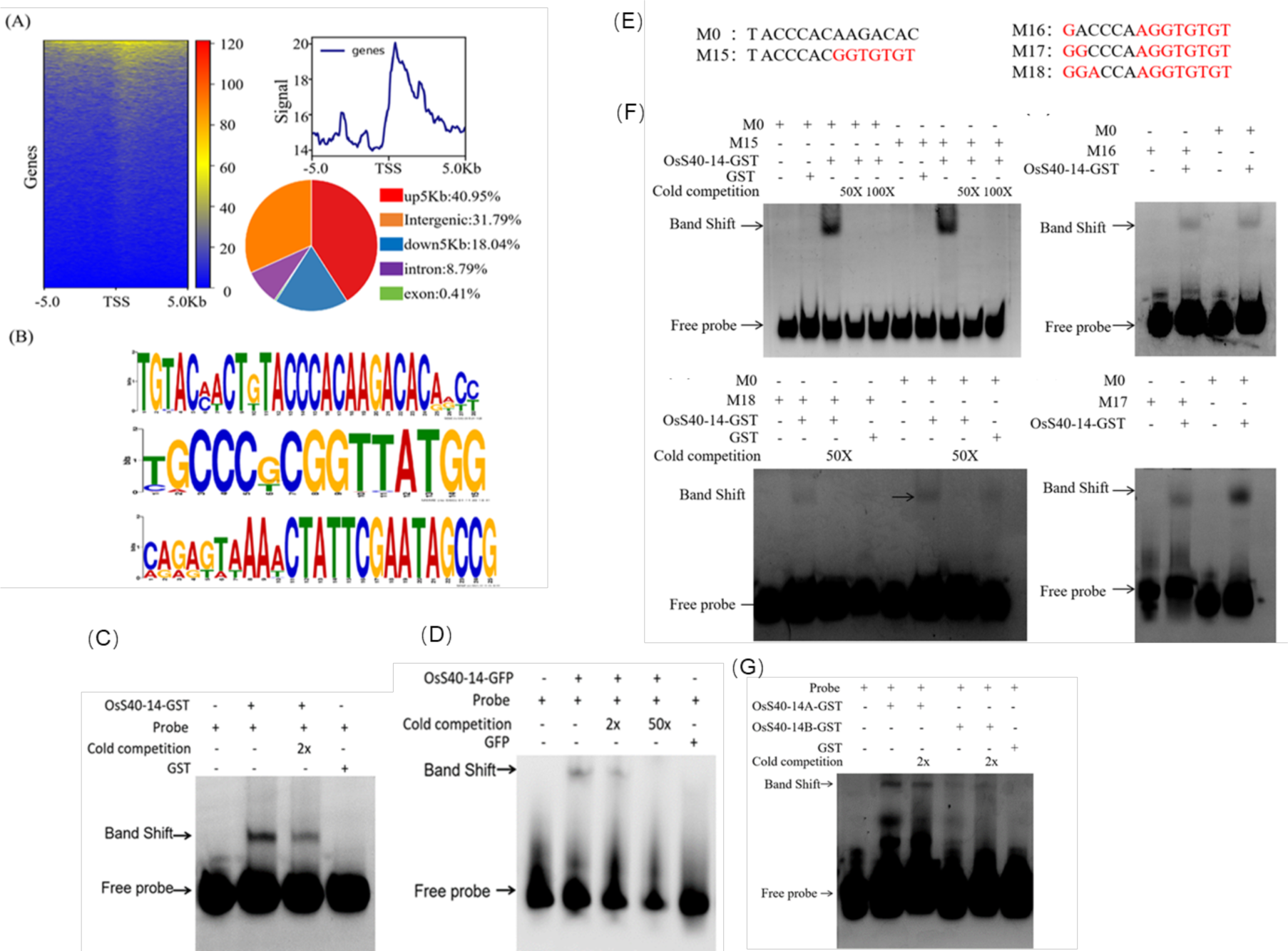
CUT&Tag dataset analysis and transcription factor OsS40-14 bound seed region identification. (A) Analysis of peak signals in GFP control and OsS40-14-GFP CUT&Tag samples. (C) Distribution map of CUT&Tag signals upstream and downstream of the gene body in each sample. Scale regions ranged from 5-kb upstream of the translation start site (TSS) to 5-kb downstream of the translation terminal site (TES). (D) Analysis of OsS40-14 enrichment targeting conserved motifs in the genome by MEME program. (E) Identification of OsS40-14 transcription factor targeting motifs by EMSA assay The DNA Probe used in the experimental group was a Cy5-labeled DNA probe:TACCCACAAGACAC-Cy5, and The DNA probe used in the Cold competition control group was an unlabeled DNA probe: TACCCACAAGACAC, the Band shift is the protein and DNA probe binding complex, and the free probe is not bind to protein. (F) Identification of OsS40-14 transcription factor binding motifs expressed in rice protoplasts. The experimental group used Cy5 fluorescently labeled DNA probe:TACCCACAAGAC AC-Cy5, and cold competition control group did not use Cy5 fluorescently labeled DNA probes: TACCCACAAGACAC, the Band shift is the protein and DNA probe binding complex, and the free probe is not bind to protein. (G) The multi-base substitution strategy to seek the DNA core motif (Seed region) for OsS40-14 transcription factor. Schematic diagram of sequential base mutations; Identification of different types of serial mutation probes binding to OsS40-14-GST; The experimental group used Cy5 fluorescently labeled DNA probes, while the cold competition control group did not use Cy5 fluorescently labeled DNA probes, the Band shift was the protein and DNA probe binding complex, and the free probe was not bind to protein. (H) Identification of the ability for two OsS40-14 deletion mutant proteins with different fragments to target DNA motifs. The experimental group used Cy5 fluorescently labeled DNA probes, while the cold competition control group did not use Cy5 fluorescently labeled DNA probes; The Band shift was a protein and DNA probe binding complex, free probe was not bind to protein.

In order to confirm the conserve sequences of OsS40-14 targeting the genome, the electrophoretic mobility shift assay (EMSA) was performed *in vitro* to confirm the interaction between OsS40-14 protein and its putative targeting elements. The recombinant OsS40-14-GST protein expressed in *E. coli* and purified with GST-bead (Supplementary Figure S6A-B), incubated with the artificial synthesized Cy5 labeled DNA probe (Supplementary Figure S6C-D) and series of mutated Cy5 labeled DNA probes, non-labeled DNA probe was used a competitor. The results of EMSA showed that the shifted complex band appeared in the lane loaded with OsS40-14 protein plus DNA probe, when added non-labeled competitor probe, the signal intensity is declined. Further, the OsS40-14 protein from rice cell (Supplementary Figure S6E-F) that transient expressed in protoplasts of rice leaf replace the OsS40-14 recombinant protein expressed in *E. coli*, the result of the EMSA is the same, indicating that OsS40-14 recombinant protein can specifically bind to the DNA fragment TACCCACAAGACAC of the genome (Figure 6E). In addition, a series of single or triple nucleotide mutated DNA probes were used to screen the core-binding motif of OsS40-14 (Figure 6F; Supplementary Figure S7), the result exhibited that ACCCA is the core-binding motif. Furthermore, the result of EMSA with various domain of OsS40-14 peptide and the DNA fragment TACCCACAAGACAC showed that the C-terminal fragment of OsS40-14 protein including NLS and DUF548 domain was DNA binding domain (Figure 6G; Supplementary Figure S6G). Therefore, OsS40-14 can specifically bind to the conserve sequence TACCCACAAGACAC of the genome, The ACCCA is the seed region of OsS40-14 targeting to genome.

### Integrative analysis of CUT&Tag-seq and RNA-seq DEGs reveals direct targets of OsS40-14 related to nutrient recycling and phosphorylation, and in responsive to ROS

To explore the potential function of OsS40-14, integrative analysis of CUT&Tag dataset and RNA-seq DEGs showed that 153 target genes of OsS40-14 were identified when overlapping the targets of CUT&Tag seq and DEGs of RNA-Seq at 0DDI (Figure 7A; Supplementary Table S10). GO enrichment analysis of 153 bound DEGs (TAGs) showed that the genes repressed TAGs by OsS40-14 were enriched mainly in chloroplast organization process and photosynthesis categories (Figure 7B). Among them, 12 transcription factors were included (Figure 7C), for example, two senescence induced transcription factor *OsNAC1* and *OsWRKY53* decreased their transcript level 3.0 and 1.5 times in the *oss40-14* mutant relative to WT, respectively (Figure 7C). All 153 TAGs including 92 upregulated and 61 downregulated target genes were categorized to 5 clusters (Figure 7D, Supplementary Table S11-12).

**Figure 7.**
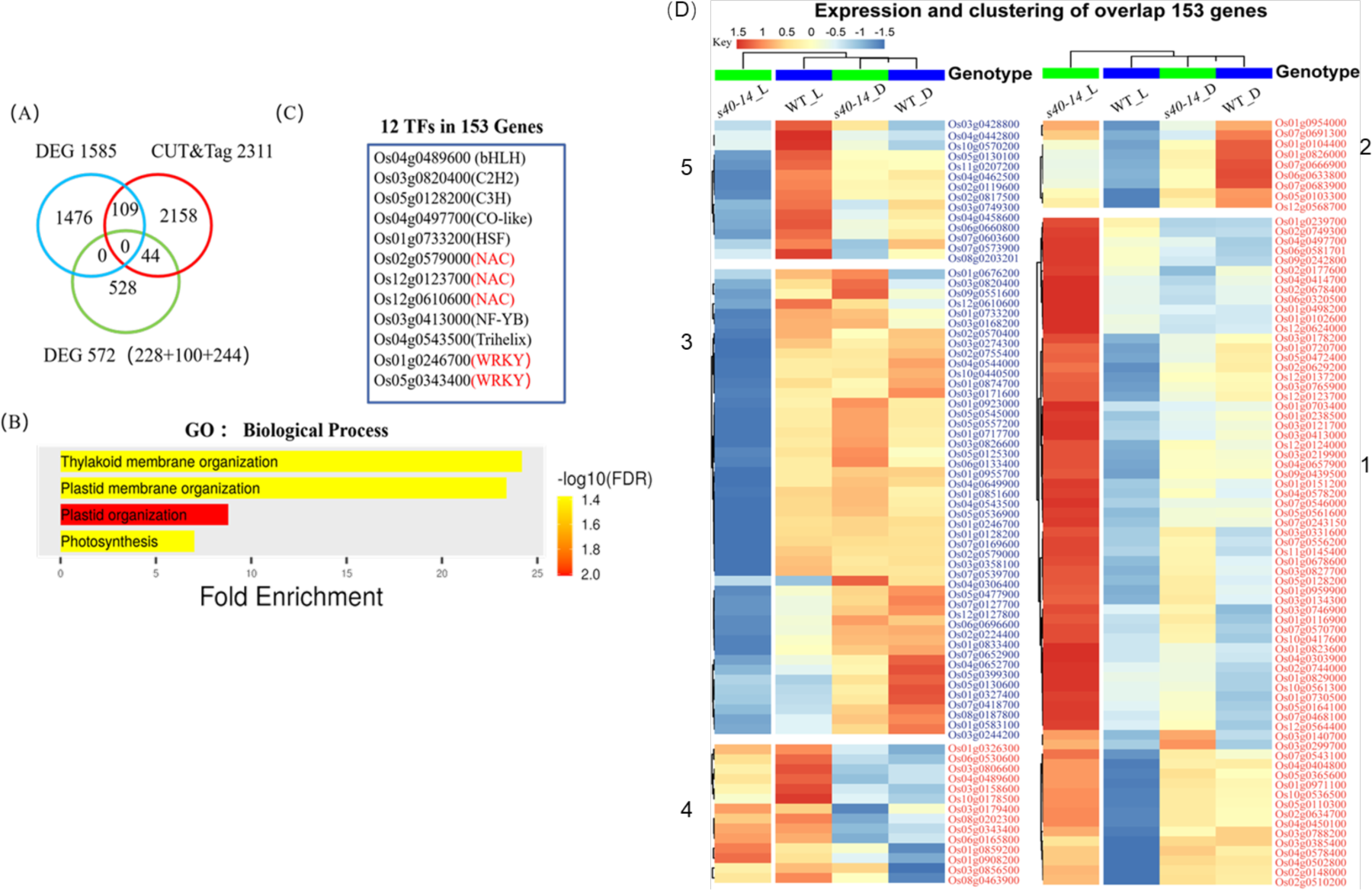
The direct targets of OsS40-14 are identified by integrative analysis of CUT&TAG-seq and RNA-seq data under dark-induced condition. (A) Venn diagram showing the overlapping target genes of the CUT&Tag-seq target gene set and the RNA-seq differentially expressed gene set; (B) GO enrichment analysis of 153 overlapping target genes; (C) comparison of 153 overlapping target genes overlapped with the known gene set of the rice genome-wide transcription factor database, which yielded a total of 12 transcription factors. (D) Expression changes of 153 candidate downstream target genes of OsS40-14 screened by joint analysis in four groups of materials. Blue font marks genes that are down-regulated in mutant leaves, while red indicates up-regulated genes. Genes with similar gene expression trends were clustered according to k=5. Heatmaps were produced using the R package heatmap program.

Among them, 7/153 dark-induced target genes (Di-TAGs) of OsS40-14 in dark-induced condition were identified by ChIP-qPCR and RT-qPCR. Two genes (*OsCHITINASE 17*/ *CHITINASE 2*, β-D-xylosidase/ Os11g0297800) encoding cell wall catabolic enzymes and four anion transporter genes (Na^+^/H^+^ ANTIPORTER/*OsNHX1*, Glucose-6-phosphate translocator 1/*OsGPT1*, Phosphate translocator 19/*OsGPT19*, Transferase family protein/ *Os06g0145600*) were selected to confirm CUT&TAG seq data. To confirm whether these genes are direct targets of OsS40-14, we performed IGV data visualization of these gene using OsS40-14-GFP targeted peak file and GFP file as the control. Interestingly, only the gene encoding the transferase family protein showed OsS40-14-GFP binding signals in its promoter region, OsGPT1 and OsGPT19 existed special CUT&Tag peak signals from OsS40-14 in both their promoter and exon regions, whereas the other 3 genes possessed OsS40-14 targeting signals mainly in their intron or exon regions (Figure 8A). By using RT-qPCR confirmation, all these seven genes were significantly upregulated under darkness treatment (Figure 8B) in the *oss40-14* mutant relative to WT. In addition, three ROS related TAGs such as ROS-producing 2OG-Fe oxygenase gene (Han et al., 2023) and GLUTATHIONE S-TRANSFERASE 46 gene (Sharma et al., 2014) both were downregulated by 1.9 times, and 2-OXOGLUTARATE-DEPENDENT DIOXYGENASE (Hu et al., 2019; Wang et al., 2022) displayed a substantial 4.6-fold decrease in the *oss40-14* mutant compared to the wild type.

**Figure 8.**
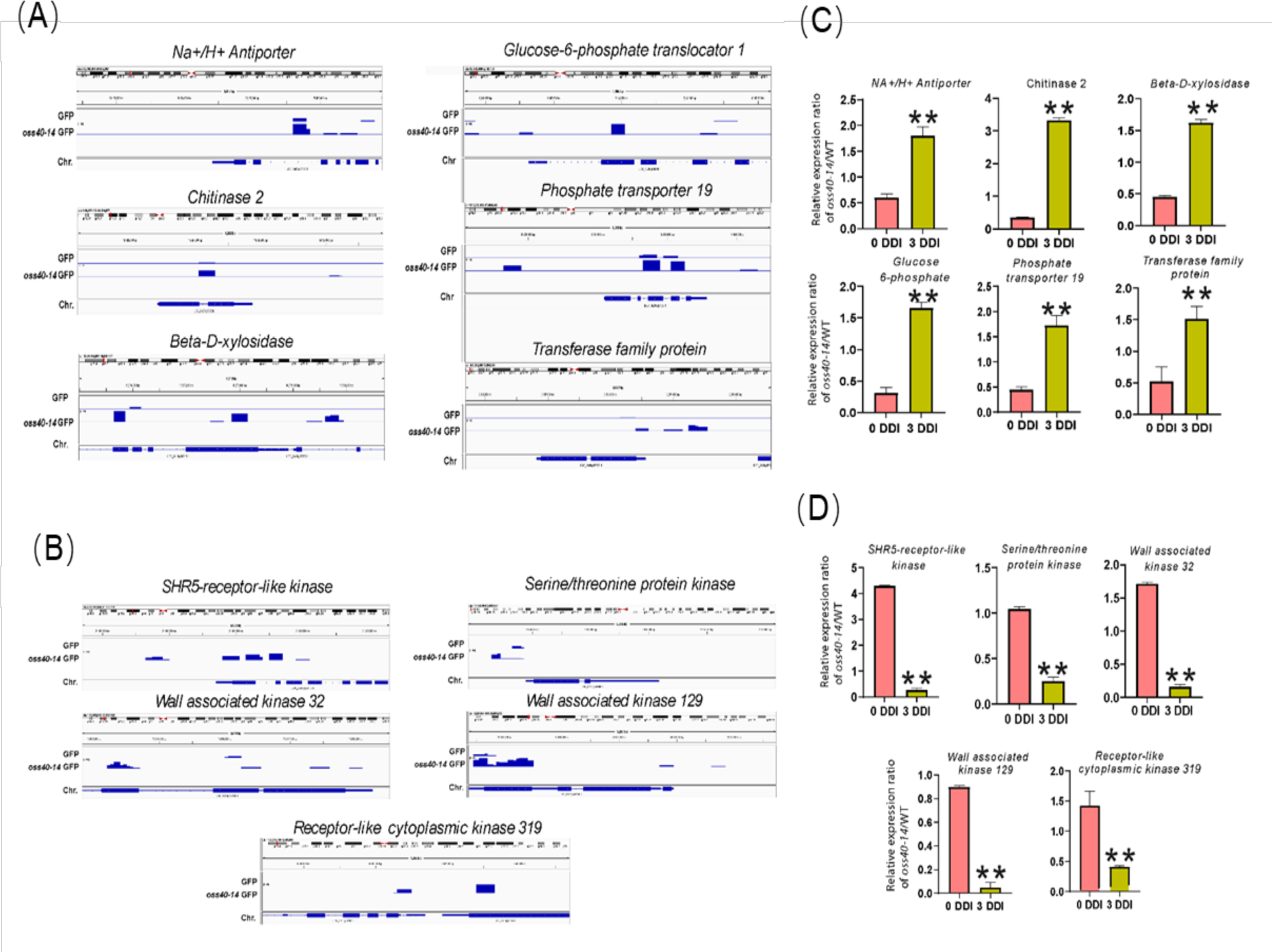
ChIP-qPCR validation of transcription factor OsS40-14 binding at the promoters of downstream genes. (A) ChIP-qPCR validation of transcription factor OsS40-14 binding at the promoters of downstream genes OsWRKY46 and OsWRKY53. The left side of (A) and (B) shows IGV visualization plots, data range=200, red peaks indicate binding positions; the right side of (A) and (B) shows the analysis of the results of ChIP-qPCR experiments, N for negative control; T-test analysis was performed using GraphPad Prism 9, with asterisks indicating significance (** indicates P<0.01), of % input normalized to GFP. (B) IGV view of 7 selected upregulated targeted genes (*Na^+^/H^+^ ANTIPORTER/OsNHX1*, *phosphate translocator 1*, *phosphate translocator 19*, *phosphate transporter 9*, *Transferase family gene*, *CHITINASE2* and *beta-D-xylosidase*) and 5 downregulated targeted genes (*WALL-ASSOCIATED KINASE GENE 32*, *WALL-ASSOCIATED KINASE GENE 129*, *SHR5-receptor-like kinase*, *serine/threonine-protein kinase, RECEPTOR-LIKE CYTOPLASMIC KINASE 319*). (C) Further Real-Time RT-PCR (qRT-PCR) of these target genes confirms the validation of DEGs in response to dark-induced leaf senescence. The endogenous *OsACTIN* gene (LOC_Os03g50885) was used as the reference gene. Asterisks indicate significant difference compared with the expression level of *OsS40-14* at 0 DDI (B, E) (Student’s t-test, *P<0.05, ***P<0.01) (A, D) DDI, day(s) of dark incubation.

The downregulated Di-TAGs were mainly enriched in protein kinase genes like WALL-ASSOCIATED KINASE GENE 32 (*OsWAK32)*, WALL-ASSOCIATED KINASE GENE 129 (*OsWAK129)*, serine/threonine-protein kinase (Os01g0137700) and RECEPTOR-LIKE CYTOPLASMIC KINASE 319 (*OsRLCK319)* in the *oss40-14* relative to WT. The expression levels of these genes were evidently increased during the dark-induced leaf senescence (Figure 8C). The IGV data visualization of these genes confirmed that the 4 genes were targeted by OsS40-14-GFP at their promoter regions, whereas RECEPTOR-LIKE CYTOPLASMIC KINASE 319 was targeted in the exon region (Figure 8D). All these five genes were significantly downregulated under darkness treatment in the *oss40-14* mutant relative to WT (Figure 8E), suggesting OsS40-14 promotes these five kinase gene expression under dark-induced senescence. Furthermore, we validated the bound DEGs of OsS40-14 by ChIP-qPCR. The results of ChIP-qPCR were basically consistent with CUT&TAG sequencing (Figure 8F). These data confirmed the notion that OsS40-14 had a profound functional involvement in protein kinase activity, cell wall catabolic process and anion transport activities during dark induced leaf senescence in rice.

Taken together, OsS40-14 may directly activate the transcript levels of phosphorylation-related genes and ROS-related genes such as catalase and APX4 genes under normal growth condition, suppress genes associated with chloroplast organization under normal growth condition. After dark induction, OsS40-14 enhances the activation of genes related to protein phosphorylation such as kinases, the repression of genes related to cell wall macromolecule catabolic process and to nutrient recycling as well as to chloroplast reorganization, resulting in dark-induced leaf senescence in rice.

## Discussion

OsS40-14 is one of members of OsS40 family, playing crucial role in dark-induced leaf senescence in rice (Habiba et al., 2021; Zheng et al., 2019). In this study, integrative analysis of RNA-seq and Cut&Tag seq data provides an overview of OsS40-14 downstream targets that either directly or indirectly affected the transcript levels during dark induced senescence condition. Under dark-induced senescence condition, genome wide transcriptome and Cut&Tag analysis revealed that OsS40-14 directly downregulated cell wall catabolic genes like OsCHITINASE 17, putative expressed beta-D-xylosidase, and anion transporter and osmotic stress related genes, such as NA+/H+ ANTIPORTER (*OsNHX1)*, PHOSPHATE TRANSLOCATOR 1 (*OsGPT1)*, PHOSPHATE TRANSLOCATOR 19/ *OsGPT19* (Figure 8C, D), while it directly upregulates various protein kinase genes such as RECEPTOR-LIKE CYTOPLASMIC KINASE 319, WALL-ASSOCIATED KINASE GENE 32, WALL-ASSOCIATED KINASE GENE 129, and serine/threonine-protein kinas/Os01g0137700 (Figure 8). OsS40-14 additionally increases ROS production to promote senescence under normal growth condition. In addition, the GO terms for upregulated non-targeted DEGs were mainly related to photosynthesis and chlorophyll biosynthesis process, and small molecule metabolic process in the *oss40-14* relative to WT (Supplementary Table S1). The GO terms for non-targeted and downregulated DEGs were mainly related to protein phosphorylation, lipid metabolism, rhythmic process, chlorophyll catabolic process, mitochondrial calcium ion transport process and response to biotic stimulus, which consistence with the phenotype of *oss40-14* and oeOsS40-14 (Figure 1-3; Habiba et al., 2021). Our finding proposed a down-stream chloroplast organization related regulatory network of OsS40-14 under dark induced senescence condition, which well explains dark induced delayed senescence phenotype in the *oss40-14* (Figure 9).

**Figure 9.**
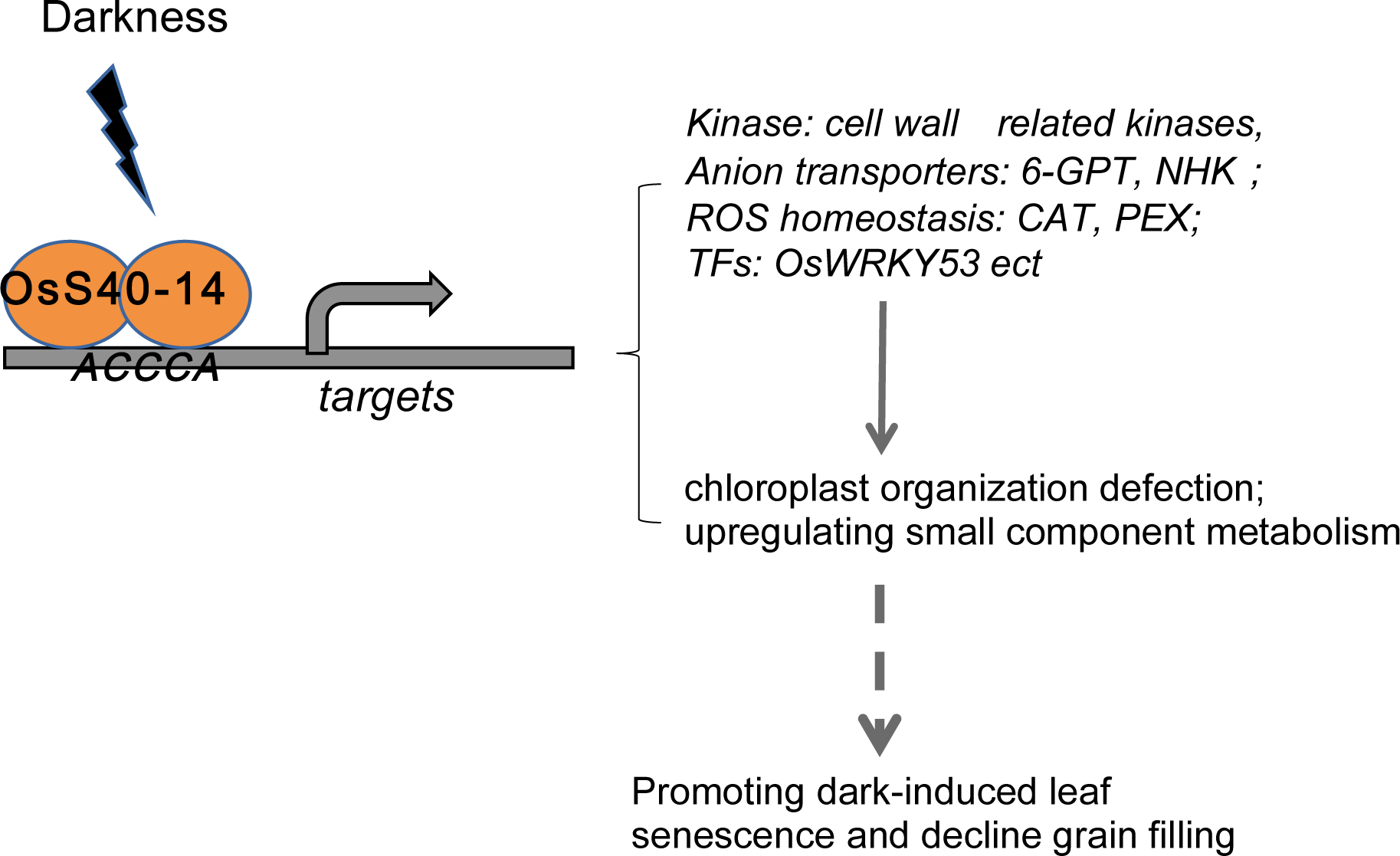
Working model of OsS40-14 regulates downstream targets during dark-induced leaf senescence in rice. The combination CUT&Tag with protoplast transient transformation system, with EMSA confirmation reveal OsS40-14 directly targeted conserve “ACCCA” seed region of downstream genes, in which OsS40-14 involved in the regulation of various biological processes related to cell wall metabolism, nutrient recycling via phosphate transporter and osmotic balance via anion transporter, as well as by promoting protein phosphorylation via kinases and ROS production to accelerate chloroplast organization defection during dark-induced leaf senescence.

Due to the unavailability of suitable antibody against OsS40-14 and instability of OsS40-14 protein, it is difficult for us to utilize ChIP-seq method and the overexpressing lines to study the binding sites of OsS40-14 in the rice genome. CUT&Tag is an enzyme-tethering method based on in situ chromatin tag mentation and used mainly for efficient profiling of epigenetic modification states in cultured animal cells (Kaya□Okur et al., 2019; 2020). Recently, CUT&Tag protocol has also been applied in plant tissues or cells for epigenomic analysis or determining binding landscape of transcription factors (Ouyang et al., 2022; Wu et al., 2022; Zhang et al., 2023). Thus, we developed the CUT&Tag manner combined with protoplast-based transient expression system for constructing the OsS40-14-bound DNA sequencing library and identified three targeted genes by ChIP-qPCR as well as 2311 potential bound genes according to the obtained CUT&Tag-seq data (Figure 6, Supplementary Figure S4). Furthermore, we used EMSA assay to identify OsS40-14 protein binding conserve seed region “ACCCA” (Figure 6). Despite some limitations in this method, like stress conditions during protoplast preparation, protoplasts mainly generated from shoots of seedlings, requirement of high transformation efficiency of plasmids, protoplast-based CUT&Tag provides a time-saving, low-cost and high-throughput approach for studying the genome-wide binding sites of plant transcription factors or DNA-related nuclear proteins, which lack suitable antibodies or stable transgenic materials. Cut&Tag assay was performed using transient transformed OsS40-14 protoplast, reflecting a transient status. In fact, OsS40-14 protein unstably existent in the tissue, its function might be transiently inducible. Therefore, current results are convincing and significant.

It has been reported that Arabidopsis cell wall glycosyl hydrolases β-glucosidase (At3g60140), defined as dark-inducible gene 2 (*DIN2*) (Fujiki et al., 2001), as well as another two glycosyl hydrolase β-xylosidase (At5g49360) (Lee et al., 2007) and β-galactosidase (At5g56870) increased in the cell wall fraction when plants were in darkness. Another enzyme related to cell wall metabolism, chitinase, is one of the important glycosyl hydrolases enzyme family that has been found to engage in stress-induced leaf senescence. Plant chitinases have effects in a number of abiotic stress conditions as well as disease resistance in normal plant growth and development (Punja and Zhang, 1993). More interestingly, one of the biofuel plant *Jatropha circuss* overexpressing Na/H antiporter (*SbNHX1*) showed better response to leaf senescence as well as salt tolerance (Jha et al., 2013). In addition, nutrient remobilization from senescing leaves to developing tissues is important, so that precious nutrients, such as phosphorus and carbohydrates, are not lost to the environment upon abscission of the fully-senescent leaf. Three phosphate transporter genes OsPT5, OsPT19 and OsPT20, are upregulated during dark stress condition (Ye et al., 2015; Wei et al., 2022) and increased expression of these genes in flag leaf contributes to remobilization of phosphorus from senescing leaves to developing grains (Jeong et al., 2017). Another phosphate translocator glucose 6-phosphate (GPT) which is a plastid translocator gene importing glucose-6-phosphate (Glc6P) into plastids for starch synthesis. Rice GPT play important roles in coordinating starch metabolism in the plastid with carbohydrate metabolism in the cytosol (Toyota et al., 2006) during leaf senescence. These examples can support our findings. Under dark induced senescence condition, OsS40-14 directly targets and down-regulates cell wall catabolic genes like *OsCHITINASE 17,* putative expressed *beta-D-xylosidase*, and *anion transporter* as well as osmotic stress related genes, such as NA+/H+ANTIPORTER (*OsNHX1*), PHOSPHATE TRANSLOCATOR 1 (*OsGPT1*), PHOSPHATE TRANSLOCATOR 19 (*OsGPT19*) (Figure 8)

Furthermore, understanding the function of several protein kinases in leaf senescence has advanced remarkably. For example, pathogen resistance, heavy-metal induced senescence and plant developmental senescence are significantly influenced by the wall-associated kinase (WAK) gene family, one of the receptor-like kinase (RLK) gene families in plants (Lally et al., 2001; Wagner and Kohorn, 2001; Sivaguru et al., 2003; Zhang et al., 2005); Arabidopsis AtWAKL10 is activated by ABA, JA, and SA, and it negatively controls leaf senescence progression; *atwakl10* mutants exhibit accelerated leaf senescence, while *AtWAKL10* overexpression plants exhibit the reverse phenotype (Li et al., 2021). However, in this study, various protein kinase genes such as RECEPTOR-LIKE CYTOPLASMIC KINASE 319, WALL-ASSOCIATED KINASE GENE 32, WALL-ASSOCIATED KINASE GENE 129 were directly downregulated under dark-induced condition in the *oss40-14* mutant, exhibiting a stay-green phenotype (Figure 8E, F), seemly inconsistence with the case of WAKL10 in *Arabidopsis*. Our findings proposed a possible mechanism of OsS40-14 under dark induced senescence condition that OsS40-14 accelerating dark-induced senescence results from a repression in the mechanical integrity of the component catabolism and nutrient recycling from the leaves to the growing section *via* a transporter but promote kinase activity and their phosphorylation signal transduction and ROS production affecting plastid organization. The detail regulatory mechanism must be further investigated.

In summary, the combination CUT&Tag with protoplast transient transformation system, with EMSA confirmation reveal OsS40-14 directly targeted conserve “ACCCA” seed region of downstream genes, in which OsS40-14 involved in the regulation of various biological processes related to cell wall metabolism, nutrient recycling *via* phosphate transporter and osmotic balance via anion transporter, as well as by promoting protein phosphorylation *via* kinases and ROS production to accelerate chloroplast organization defection during dark-induced leaf senescence.

### Gene information

Sequence data of genes from this article can be found in the National Center for Biotechnology Information (NCBI): *OsS40-14* (*Os05g0531000*, OSNPB_050148000); *OsS40-1* (*Os05g0531100*, OSNPB_050531100); *OsS40-2* (*Os05g0518800*, OSNPB_050518800); *OsS40-12* (*Os11g0154300*, OSNPB_110154300); *UBQ* (*Os01g0328400*, OSNPB_010328400); *OsNHX1* (*Os07g0666900*, OSNPB_070666900); *Phosphate Translocator 1* (*Os08g0187800*, OSNPB_080187800); *Phosphate Transporter 19* (*Os09g0454600*, OSNPB_090454600); *beta-D-xylosidase* (*Os11g0297800*, OSNPB_110297800); transferase family protein (*Os06g0145600*, OSNPB_060145600); *SHR5-receptor-like kinase* (*Os08g0202300*, OSNPB_080202300); *WALL-ASSOCIATED KINASE GENE 32* (*Os04g0307500*, OSNPB_040307500); *serine/threonine-protein kinase* (*Os01g0137700*); *WALL-ASSOCIATED KINASE GENE 129* (*Os12g0615300*, OSNPB_120615300); *RECEPTOR-LIKE CYTOPLASMIC KINASE 319* (*Os11g0225000*, OSNPB_110225000).

### Supplementary data

Fig S1 Expression of OsS40-1, 2, and 12 measured in 10-days-old rice seedling after complete darkness treatment for 0DDI to 6 DDI at 28 °C.

Fig S2 Generation and Identification of oss40-14 knockout mutant using the CRISPR/Cas9 system.

Fig S3 OsS40-14 overexpressing transgenic rice plants show a stay-green phenotype during dark-induced leaf senescence.

Fig S4 DEG profile of *oss40-14* versus wild-type (ZH11) ex vivo leaves after 3 days of dark treatment.

Fig S5 Detection of the library of CUT&Tag samples.

Fig S6 Recombinant OsS40-14 protein, OsS40-14 mutated protein, OsS40-14 protein isolated from transformed protoplast and were detected

Fig S7 Cy5 fluorescently labeled DNA probes were prepared by LUEGO short linker complementary method

Fig S8 The single-base substitution substitution strategy to seek the DNA core motif for OsS40-14 transcription factor

Fig S9 GO Enrichment of OsS40-14 targets of a CUT&Tag data. Table S1. Differentially expressed genes in venn recult 4 treatment sets

Table S2. Differentially expressed genes in *oss40-14* mutant compare to WT at ODDI. Table S3. Differentially expressed genes of 3DDI vs 0DDI in the WT.

Table S4. Differentially expressed genes of 3DDI vs 0DDI in oss40-14 mutant

Table S5. Differentially expressed genes in *oss40-14* mutant compare to WT at 3DDI. Table S6. 74 Transcription factors venn rice TFs vs DEG_all.

Table S7. DEGs related to hormones in *oss40-14* mutant compare to WT at 3DDI relative to 0DDI.

Table S8 45 SAG venn Rice SAGs 225 vs DEG_all. Table S9. CUT-Tag peaks of OsS40-14-GFP and GFP. Table S10. THE ordered ID Description_153 genes.

Table S11. 153_overlap_cluster. Table S12. 12 TFs in 153 cluster.

Table S13. The list of primer sequences used in this study.

## AUTHOR CONTRIBUTIONS

X.Z and Y.M conceived and designed the research. H., C.F., W.H., Y.S., X.W., W.W., and Y.L collected the data. H., C.F., W.L., and N.A analyzed the data. H., X.Z, and Y.M draft and revised the manuscript. All authors read and approved the final manuscript.

## Data availability

RNA-seq raw data are available from the Sequence Read Archive (https://www.ncbi.nlm.nih.gov/sra) under accession number PRJNA1135482. All data generated or analyzed during this study are included in this published article and its supplementary information files.

## Reference

1. Adabnejad H, Kavousi HR, Hamidi H, Tavassolian I. 2015. Assessment of the vacuolar Na + /H + antiporter (NHX1) transcriptional changes in Leptochloa fusca L. in response to salt and cadmium stresses. Molecular Biology Research Communications 4, 133–142.

2. Ao Y, Li Z, Feng D, Xiong F, Liu J, Li JF, Wang M, Wang J, Liu B, Wang H Bin. 2014. OsCERK1 and OsRLCK176 play important roles in peptidoglycan and chitin signaling in rice innate immunity. Plant Journal 80, 1072–1084.

3. Apse MP, Aharon GS, Snedden WA, Blumwald E. 1999. Salt tolerance conferred by overexpression of a vacuolar Na+/H+ antiport in Arabidopsis. Science 285, 1256–1258.

4. Balazadeh S, Kwasniewski M, Caldana C, Mehrnia M, Zanor MI, Xue GP, Mueller-Roeber B. 2011. ORS1, an H2O2-responsive NAC transcription factor, controls senescence in arabidopsis thaliana. Molecular Plant 4, 346–360.

5. Becker W, Apel K. 1993. Differences in gene expression between natural and artificially induced leaf senescence. Planta 189, 74–79.

6. Blumwald E, Poole RJ. 1986. Kinetics of Ca 2+ /H + Antiport in Isolated Tonoplast Vesicles from Storage Tissue of Beta vulgaris L. . Plant Physiology 80, 727–731.

7. Breeze E, Harrison E, McHattie S, et al. 2011. High-resolution temporal profiling of transcripts during Arabidopsis leaf senescence reveals a distinct chronology of processes and regulation. Plant Cell 23, 873–894.

8. Brouwer B, Ziolkowska A, Bagard M, Keech O, Gardeström P. 2012. The impact of light intensity on shade-induced leaf senescence. Plant, Cell and Environment 35, 1084–1098.

9. Buchanan-Wollaston V, Earl S, Harrison E, Mathas E, Navabpour S, Page T, Pink D. 2002. The molecular analysis of leaf senescence - a genomics approach. Plant Biotechnology Journal 1, 3–22.

10. Buchanan-Wollaston V, Page T, Harrison E, et al. 2005. Comparative transcriptome analysis reveals significant differences in gene expression and signalling pathways between developmental and dark/starvation-induced senescence in Arabidopsis. Plant Journal 42, 567– 585.

11. Cao J, Zhang Y, Tan S, Yang Q, Wang H-L, Xia X, Luo J, Guo H, Zhang Z, Li Z. 2022. LSD 4.0: an improved database for comparative studies of leaf senescence. Molecular Horticulture 2.

12. Castillo MC, Lozano-Juste J, González-Guzmán M, Rodriguez L, Rodriguez PL, León J. 2015. Inactivation of PYR/PYL/RCAR ABA receptors by tyrosine nitration may enable rapid inhibition of ABA signaling by nitric oxide in plants. Science Signaling 8.

13. Chen H, An R, Tang JH, Cui XH, Hao FS, Chen J, Wang XC. 2007. Over-expression of a vacuolar Na+/H+ antiporter gene improves salt tolerance in an upland rice. Molecular Breeding 19, 215–225.

14. Chen LQ, Hou BH, Lalonde S, et al. 2010. Sugar transporters for intercellular exchange and nutrition of pathogens. Nature 468, 527–532.

15. Chen L, Jameson GB, Guo Y, Song J, Jameson PE. 2022. The LONELY GUY gene family: from mosses to wheat, the key to the formation of active cytokinins in plants. Plant Biotechnology Journal 20, 625–645.

16. Chen W ping, Kuo T teh. 1993. A simple and rapid method for the preparation of gram-negative bacterial genomic DNA. Nucleic Acids Research 21, 2260.

17. Dietrich K, Weltmeier F, Ehlert A, Weiste C, Stahl M, Harter K, Dröge-Lasera W. 2011. Heterodimers of the Arabidopsis transcription factors bZIP1 and bZIP53 reprogram amino acid metabolism during Low energy stress. Plant Cell 23, 381–395.

18. Fan J, Bai P, Ning Y, et al. 2018. The Monocot-Specific Receptor-like Kinase SDS2 Controls Cell Death and Immunity in Rice. Cell Host and Microbe 23, 498–510.e5.

19. Fatima M, Ma X, Zhou P, Zaynab M, Ming R. 2021. Auxin regulated metabolic changes underlying sepal retention and development after pollination in spinach. BMC Plant Biology 21, 1–15.

20. Fujiki Y, Yoshikawa Y, Sato T, Inada N, Ito M, Nishida I, Watanabe A. 2001. Dark-inducible genes from Arabidopsis thaliana are associated with leaf senescence and repressed by sugars. Physiologia Plantarum 111, 345–352.

21. Fukuda A, Nakamura A, Tagiri A, Tanaka H, Miyao A, Hirochika H, Tanaka Y. 2004. Function, Intracellular Localization and the Importance in Salt Tolerance of a Vacuolar Na+/H+ Antiporter from Rice. Plant and Cell Physiology 45, 146–159.

22. Gad AG, Habiba, Zheng X, Miao Y. 2021. Low light/darkness as stressors of multifactor-induced senescence in rice plants. MDPI AG.

23. Ge L, Chen H, Jiang JF, Zhao Y, Xu ML, Xu YY, Tan KH, Xu ZH, Chong K. 2004. Overexpression of OsRAA1 causes pleiotropic phenotypes in transgenic rice plants, including altered leaf, flower, and root development and root response to gravity. Plant Physiology 135, 1502–1513.

24. Van Der Graaff E, Schwacke R, Schneider A, Desimone M, Flügge UI, Kunze R. 2006. Transcription analysis of arabidopsis membrane transporters and hormone pathways during developmental and induced leaf senescence. Plant Physiology 141, 776–792.

25. Guo Y. 2013. Towards systems biological understanding of leaf senescence. Plant Molecular Biology 82, 519–528.

26. Guo Y, Gan S. 2006. AtNAP, a NAC family transcription factor, has an important role in leaf senescence. Plant Journal 46, 601–612.

27. Guo Y, Gan SS. 2012. Convergence and divergence in gene expression profiles induced by leaf senescence and 27 senescence-promoting hormonal, pathological and environmental stress treatments. Plant, Cell and Environment 35, 644–655.

28. Guo P, Li Z, Huang P, Li B, Fang S, Chu J, Guo H. 2017. A tripartite amplification loop involving the transcription factor WRKY75, salicylic acid, and reactive oxygen species accelerates leaf senescence. Plant Cell 29, 2854–2870.

29. Guo Y, Ren G, Zhang K, Li Z, Miao Y, Guo H. 2021. Leaf senescence: progression, regulation, and application. Molecular Horticulture 1.

30. Habiba, Xu J, Gad AG, et al. 2021. Five OsS40 Family Members Are Identified as Senescence-Related Genes in Rice by Reverse Genetics Approach. Frontiers in Plant Science 12.

31. Han J, Wang X, Niu S. 2023. Genome-Wide Identification of 2-Oxoglutarate and Fe (II)-Dependent Dioxygenase (2ODD-C) Family Genes and Expression Profiles under Different Abiotic Stresses in Camellia sinensis (L.). Plants 12.

32. Hasegawa M, Bressan R, Pardo JM. 2000. The dawn of plant salt tolerance genetics. Trends in Plant Science 5, 317–319.

33. Hayashi K and Kojima C, 2008, pCold-GST vector: A novel cold-shock vector containing GST tag for soluble protein production. Protein Expression and Purification 62(1): 120–127

34. He F, Zhang F, Sun W, Ning Y, Wang GL. 2018. A Versatile Vector Toolkit for Functional Analysis of Rice Genes. Rice 11.

35. Hu T, Wang Y, Wang Q, Dang N, Wang L, Liu C, Zhu J, Zhan X. 2019. The tomato 2-oxoglutarate-dependent dioxygenase gene SlF3HL is critical for chilling stress tolerance. Horticulture Research 6.

36. Huang J, Cai M, Long Q, Liu L, Lin Q, Jiang L, Chen S, Wan J. 2014. OsLOX2, a rice type I lipoxygenase, confers opposite effects on seed germination and longevity. Transgenic Research 23, 643–655.

37. Jeong K, Baten A, Waters DLE, Pantoja O, Julia CC, Wissuwa M, Heuer S, Kretzschmar T, Rose TJ. 2017. Phosphorus remobilization from rice flag leaves during grain filling: an RNA1seq study. Plant Biotechnol J. 2017 Jan; 15(1): 15–26.

38. Jha B, Mishra A, Jha A, Joshi M. 2013. Developing Transgenic Jatropha Using the SbNHX1 Gene from an Extreme Halophyte for Cultivation in Saline Wasteland. PLoS ONE 8.

39. Jin Y, Pan W, Zheng X, Cheng X, Liu M, Ma H, Ge X. 2018. OsERF101, an ERF family transcription factor, regulates drought stress response in reproductive tissues. Plant Molecular Biology 98, 51–65.

40. Kanazawa S, Sano S, Koshiba T, Ushimaru T. 2000. Changes in antioxidative enzymes in cucumber cotyledons during natural senescence: Comparison with those during dark-induced senescence. Physiologia Plantarum 109, 211–216.

41. Karimi M, Inzé D, Depicker A. 2002. GATEWAY vectors for Agrobacterium-mediated plant transformation. Trends in Plant Science 7, 193–195.

42. Kaya□Okur HS, Janssens DH, Henikoff JG, Ahmad K, and Henikoff S. 2020. Efficient low1 cost chromatin profiling with CUT&Tag. Nat. Protoc. 15: 3264–3283

43. Kaya□Okur HS, Wu SJ, Codomo CA, Pledger ES, Bryson TD, Henikoff JG, Ahmad K, and Henikoff S. 2019. CUT&Tag for efficient epigenomic profiling of small samples and single cells. Nat. Commun. 10: 1930.

44. Kim H, Kim HJ, Vu QT, Jung S, Robertson McClung C, Hong S, Nam HG. 2018. Circadian control of ORE1 by PRR9 positively regulates leaf senescence in Arabidopsis. Proceedings of the National Academy of Sciences of the United States of America 115, 8448–8453.

45. Krupinska K, Haussühl K, Schäfer A, van der Kooij TA, Leckband G, Lörz H, Falk J. 2002. A novel nucleus-targeted protein is expressed in barley leaves during senescence and pathogen infection. Plant Physiol. 130(3):1172–80.

46. Kumar S, Kalita A, Srivastava R, Sahoo L. 2017. Co-expression of arabidopsis NHX1 and bar improves the tolerance to salinity, oxidative stress, and herbicide in transgenic mungbean. Frontiers in Plant Science 8.

47. Kumar K, Sinha AK. 2013. Overexpression of constitutively active mitogen activated protein kinase kinase 6 enhances tolerance to salt stress in rice. Rice 6, 1–5.

48. Kusaba M, Ito H, Morita R, et al. 2007. Rice non-yellow coloring1 is involved in light-harvesting complex II and grana degradation during leaf senescence. Plant Cell 19, 1362–1375.

49. Kusaba M, Tanaka A, Tanaka R. 2013. Stay-green plants: What do they tell us about the molecular mechanism of leaf senescence.

50. Lally D, Ingmire P, Tong HY, He ZH. 2001. Antisense expression of a cell wall-associated protein kinase, WAK4, inhibits cell elongation and alters morphology. Plant Cell 13, 1317–1331.

51. Langmead B, and Salzberg SL. 2012. Fast gapped-read alignment with Bowtie 2. Nat. Methods 9: 357–354.

52. Lee SK, Kim BG, Kwon TR, et al. 2011. Overexpression of the mitogen-activated protein kinase gene OsMAPK33 enhances sensitivity to salt stress in rice (Oryza sativa L.). https://link.springer.com/article/10.1007/s12038-011-9002-8. Accessed July 2023.

53. Lee H-N, Lee K-H, Kim CS. 2015. Abscisic acid receptor PYRABACTIN RESISTANCE-LIKE 8, PYL8, is involved in glucose response and dark-induced leaf senescence in Arabidopsis. Biochemical and Biophysical Research Communications 463, 24–28.

54. Lee EJ, Matsumura Y, Soga K, Hoson T, Koizumi N. 2007. Glycosyl hydrolases of cell wall are induced by sugar starvation in arabidopsis. Plant and Cell Physiology 48, 405–413.

55. Lee RH, Wang CH, Huang LT, Chen SG. 2001. Leaf senescence in rice plants: Cloning and characterization of senescence up-regulated genes. Journal of Experimental Botany 52, 1117–1121.

56. Li L, Li K, Ali A, Guo Y. 2021. Atwakl10, a cell wall associated receptor-like kinase, negatively regulates leaf senescence in arabidopsis thaliana. International Journal of Molecular Sciences 22.

57. Li Y, Liao S, Mei P, Pan Y, Zhang Y, Zheng X, Xie Y, and Miao Y. 2021. OsWRKY93 Dually Functions Between Leaf Senescence and in Response to Biotic Stress in Rice. Front. Plant Sci. 12:643011

58. Liang C, Wang Y, Zhu Y, et al. 2014. OsNAP connects abscisic acid and leaf senescence by fine-tuning abscisic acid biosynthesis and directly targeting senescence-associated genes in rice. Proceedings of the National Academy of Sciences of the United States of America 111, 10013–10018.

59. Lichtenthaler HK, Wellburn AR. 1983. Determinations of total carotenoids and chlorophylls a and b of leaf extracts in different solvents . Biochemical Society Transactions 11, 591–592.

60. Liebsch D, Keech O. 2016. Dark-induced leaf senescence: new insights into a complex light-dependent regulatory pathway. New Phytologist 212, 563–570.

61. Liu W., Xie X., Ma X., Li J., Chen J., and Liu Y.-G. 2015. DSDecode: A Web-based Tool for Decoding of Sequencing Chromatograms for Genotyping of Targeted Mutations. Mol. Plant 8(9):1431–1433.

62. Lou D, Wang H, Liang G, Yu D. 2017. OsSAPK2 confers abscisic acid sensitivity and tolerance to drought stress in rice. Frontiers in Plant Science 8.

63. Love MI, Anders S, Huber W. 2014. Differential analysis of count data - the DESeq2 package. Genome Biology 15, 550.

64. Lu F, and Lionnet T. 2021. Transcription Factor Dynamics. Cold Spring Harbor perspectives in biology, 13 (11), 10.1101/cshperspect.a040949

65. Lv J, Shang L, Chen Y, et al. 2020. OsSLC1 Encodes a Pentatricopeptide Repeat Protein Essential for Early Chloroplast Development and Seedling Survival. Rice 13.

66. Ma B, He SJ, Duan KX, Yin CC, Chen H, Yang C, et al. 2013. Identification of rice ethylene-response mutants and characterization of MHZ7/OsEIN2 in distinct ethylene response and yield trait regulation. Mol. Plant 6, 1830–1848.

67. Mair A, Pedrotti L, Wurzinger B, et al. 2015. SnRK1-triggered switch of bZIP63 dimerization mediates the low-energy response in plants. eLife 4.

68. Malukani KK, Ranjan A, Hota SJ, Patel HK, Sonti R V. 2020. Dual activities of receptor-like kinase OsWAKL21.2 induce immune responses. Plant Physiology 183, 1345– 1363.

69. Miao Y, Laun T, Zimmermann P, Zentgraf U. 2004. Targets of the WRKY53 transcription factor and its role during leaf senescence in Arabidopsis. Plant Molecular Biology 55, 853–867.

70. Morita R, Sato Y, Masuda Y, Nishimura M, Kusaba M. 2009. Defect in non-yellow coloring 3, an α/β hydrolase-fold family protein, causes a stay-green phenotype during leaf senescence in rice. Plant Journal 59, 940–952.

71. Ortiz-Morea FA, Liu J, Shan L, He P. 2022. Malectin-like receptor kinases as protector deities in plant immunity. Nature Plants 8, 27–37.

72. Ouyang SQ, Liu YF, Liu P, Lei G, He SJ, Ma B, Zhang WK, Zhang JS, Chen SY. 2010. Receptor-like kinase OsSIK1 improves drought and salt stress tolerance in rice (Oryza sativa) plants. Plant Journal 62, 316–329.

73. Ouyang W, Luan S, Xiang X, Guo M, Zhang Y, Li G, and Li X. 2022. Profiling plant histone modification at single1cell resolution using snCUT&Tag. Plant Biotechnol. J. 20: 420–422.

74. Paluch-Lubawa E, Stolarska E, Sobieszczuk-Nowicka E. 2021. Dark-Induced Barley Leaf Senescence – A Crop System for Studying Senescence and Autophagy Mechanisms. Frontiers in Plant Science 12.

75. Park SY, Yu JW, Park JS, et al. 2007. The senescence-induced staygreen protein regulates chlorophyll degradation. Plant Cell 19, 1649–1664.

76. Pu CX, Han YF, Zhu S, et al. 2017. The rice receptor-like kinases DWARF AND RUNTISH SPIKELET1 and 2 repress cell death and affect sugar utilization during reproductive development. Plant Cell 29, 70–89.

77. Punja ZK, Zhang YY. 1993. Plant chitinases and their roles in resistance to fungal diseases. Journal of nematology 25, 526–40.

78. Pyung OL, Hyo JK, Hong GN. 2007. Leaf senescence. Annual Review of Plant Biology 58, 115–136.

79. Qiu K, Li Z, Yang Z, et al. 2015. EIN3 and ORE1 Accelerate Degreening during Ethylene-Mediated Leaf Senescence by Directly Activating Chlorophyll Catabolic Genes in Arabidopsis. PLoS Genetics 11.

80. Ramkumar MK, Senthil Kumar S, Gaikwad K, Pandey R, Chinnusamy V, Singh NK, Singh AK, Mohapatra T, Sevanthi AM. 2019. A novel stay-green mutant of rice with delayed leaf senescence and better harvest index confers drought tolerance. Plants 8.

81. Robinson JT, Thorvaldsdóttir H, Winckler W, Guttman M, Lander ES, Getz G, Mesirov JP. 2011. Integrative Genomics Viewer. Nature Biotechnology 29, 24–26.

82. Sade N, Del Mar Rubio-Wilhelmi M, Umnajkitikorn K, Blumwald E. 2018. Stress-induced senescence and plant tolerance to abiotic stress. Journal of Experimental Botany 69, 845–853.

83. Sakuraba Y, Jeong J, Kang MY, Kim J, Paek NC, Choi G. 2014. Phytochrome-interacting transcription factors PIF4 and PIF5 induce leaf senescence in Arabidopsis. Nature Communications 5.

84. Saleh A, Alvarez-Venegas R, Avramova Z. 2008. An efficient chromatin immunoprecipitation (ChIP) protocol for studying histone modifications in Arabidopsis plants. Nature Protocols 3, 1018–1025.

85. Schmittgen TD, Livak KJ. 2008. Analyzing real-time PCR data by the comparative CT method. Nature Protocols 3, 1101–1108.

86. Shannon P, Markiel A, Ozier O, Baliga NS, Wang JT, Ramage D, Amin N, Schwikowski B, Ideker T. 2003. Cytoscape: A software Environment for integrated models of biomolecular interaction networks. Genome Research 13, 2498–2504.

87. Shao L, Shu Z, Peng CL, Lin ZF, Yang CW, Gu Q. 2008. Enhanced sensitivity of Arabidopsis anthocyanin mutants to photooxidation: A study with fluorescence imaging. Functional Plant Biology 35, 714–724.

88. Sharma R, Sahoo A, Devendran R, Jain M. 2014. Over-expression of a rice tau class glutathione S-transferase gene improves tolerance to salinity and oxidative stresses in arabidopsis. PLoS ONE 9.

89. Shim Y, Kang K, An G, Paek NC. 2019. Rice DNA-Binding One Zinc Finger 24 (OsDOF24) Delays Leaf Senescence in a Jasmonate-Mediated Pathway. Plant and Cell Physiology 60, 2065–2076.

90. Sivaguru M, Ezaki B, He ZH, Tong H, Osawa H, Baluška F, Volkmann D, Matsumoto H. 2003. Aluminum-induced gene expression and protein localization of a cell wall-associated receptor kinase in Arabidopsis. Plant Physiology 132, 2256–2266.

91. Sobieszczuk-Nowicka E, Wrzesiński T, Bagniewska-Zadworna A, Kubala S, Rucińska-Sobkowiak R, Polcyn W, Misztal L, Mattoo AK. 2018. Physio-genetic dissection of dark-induced leaf senescence and timing its reversal in Barley1[OPEN]. Plant Physiology 178, 654–671.

92. Song Y, Yang C, Gao S, Zhang W, Li L, Kuai B. 2014. Age-Triggered and Dark-Induced Leaf Senescence Require the bHLH Transcription Factors PIF3, 4, and 5. Molecular Plant 7, 1776–1787.

93. Spielmeyer W, Ellis MH, Chandler PM. 2002. Semidwarf (sd-1), ‘green revolution’ rice, contains a defective gibberellin 20-oxidase gene. Proceedings of the National Academy of Sciences of the United States of America 99, 9043–9048.

94. Tang Y, Li M, Chen Y, Wu P, Wu G, Jiang H. 2011. Knockdown of OsPAO and OsRCCR1 cause different plant death phenotypes in rice. Journal of Plant Physiology 168, 1952–1959.

95. Tian X, Wang Z, Li X, Lv T, Liu H, Wang L, Niu H, Bu Q. 2015. Characterization and Functional Analysis of Pyrabactin Resistance-Like Abscisic Acid Receptor Family in Rice. Rice 8.

96. Toyota K, Tamura M, Ohdan T, Nakamura Y. 2006. Expression profiling of starch metabolism-related plastidic translocator genes in rice. Planta 223, 248–257.

97. Verma D, Singla-Pareek SL, Rajagopal D, Reddy MK, Sopory SK. 2007. Functional validation of a novel isoform of Na+/H+ antiporter from Pennisetum glaucum for enhancing salinity tolerance in rice. Journal of Biosciences 32, 621–628.

98. Wagner TA, Kohorn BD. 2001. Wall-associated kinases are expressed throughout plant development and are required for cell expansion. Plant Cell 13, 303–318.

99. Wang Y, Huan Q, Li K, Qian W. 2021. Single-cell transcriptome atlas of the leaf and root of rice seedlings. Journal of Genetics and Genomics 48, 881–898.

100. Wang H, Liu S, Fan F, Yu Q, Zhang P. 2022. A Moss 2-Oxoglutarate/Fe(II)-Dependent Dioxygenases (2-ODD) Gene of Flavonoids Biosynthesis Positively Regulates Plants Abiotic Stress Tolerance. Frontiers in Plant Science 13, 1–19.

101. Wei X, Fu Y, Yu R, Wu L, Wu Z, Tian P, Li S, Yang X, Yang M. 2022. Comprehensive sequence and expression profile analysis of the phosphate transporter gene family in soybean. Scientific Reports 12.

102. Woo HR, Kim HJ, Lim PO, Nam HG. 2019. Leaf Senescence: Systems and Dynamics Aspects. Annual Review of Plant Biology 70, 347–376.

103. Wu L, Luo Z, Shi Y, Jiang Y, Li R, Miao X, Yang F, Li Q, Zhao H, Xue J, Xu S, Zhang T, Li L. 2022. A cost-effective tsCUT&Tag method for profiling transcription factor binding landscape. J Integr Plant Biol. 64(11):2033–2038.

104. Wu XY, Kuai BK, Jia, JZ and Jing HC. 2012. Regulation of Leaf Senescence and Crop Genetic Improvement†. Journal of Integrative Plant Biology, 54: 936–952.

105. Xie W, Li X, Wang S, Yuan M. 2022. OsWRKY53 Promotes Abscisic Acid Accumulation to Accelerate Leaf Senescence and Inhibit Seed Germination by Downregulating Abscisic Acid Catabolic Genes in Rice. Frontiers in Plant Science 12.

106. Xue ZY, Zhi DY, Xue GP, Zhang H, Zhao YX, Xia GM. 2004. Enhanced salt tolerance of transgenic wheat (Tritivum aestivum L.) expressing a vacuolar Na+/H+ antiporter gene with improved grain yields in saline soils in the field and a reduced level of leaf Na +. Plant Science 167, 849–859.

107. Yang J, Worley E, Udvardi M. 2014. A NAP-AAO3 regulatory module promotes chlorophyll degradation via aba biosynthesis in arabidopsis leavesw open. Plant Cell 26, 4862–4874.

108. Ye Y, Yuan J, Chang X, Yang M, Zhang L, Lu K, Lian X. 2015. The phosphate transporter gene OsPht1;4 is involved in phosphate homeostasis in rice. PLoS ONE 10.

109. Yoshida S, Forno DA, Cock J. 1971. Laboratory Manual for Physiological Studies of Rice. International Rice Research Institute.

110. Yu T, Cen Q, Kang L, Mou W, Zhang X, Fang Y, Zhang X, Tian Q, Xue D. 2022. Identification and expression pattern analysis of the OsSnRK2 gene family in rice. Frontiers in Plant Science 13.

111. Yuchun RAO, Ran JIAO, Sheng WANG, et al. 2021. SPL36 Encodes a Receptor-like Protein Kinase that Regulates Programmed Cell Death and Defense Responses in Rice. Rice 14.

112. Zentgraf U. 2001. Identification of a transcription factor specifically expressed at the onset of leaf senescence. Planta 213, 469–473.

113. Zhang S, Chen C, Li L, Meng L, Singh J, Jiang N, Deng XW, He ZH, Lemaux PG. 2005. Evolutionary expansion, gene structure, and expression of the rice wall-associated kinase gene family. Plant Physiology 139, 1107–1124.

114. Zhang TQ, Chen Y, Liu Y, Lin WH, Wang JW. 2021. Single-cell transcriptome atlas and chromatin accessibility landscape reveal differentiation trajectories in the rice root. Nature Communications 12.

115. Zhang Y, Liu Q, Zhang Y, Chen Y, Yu N, Cao Y, Zhan X, Cheng S, Cao L. 2019. LMM24 Encodes Receptor-Like Cytoplasmic Kinase 109, Which Regulates Cell Death and Defense Responses in Rice. mdpi.com doi: 10.3390/ijms20133243.

116. Zhang Y, Li Y, Zhang Y, Zhang Z, Zhang D, Wang X, Lai B, Huang D, Gu L, Xie Y, Miao Y. 2022. Genome-wide H3K9 acetylation level increases with age-dependent senescence of flag leaf in rice. J Exp Bot. 73(14):4696–4715.

117. Zhang Y, Liu T, Meyer CA, Eeckhoute J, Johnson DS, Bernstein BE, Nusbaum C, Myers RM, Brown M, Li W and Liu XS. 2008. Model-based analysis of ChIP-Seq (MACS). Genome Biol. 9: R137.

118. Zhang Y, Zang Y, Chen J, Feng S, Zhang Z, Hu Y, Zhang T. 2023. A truncated ETHYLENE INSENSITIVE3-like protein, GhLYI, regulates senescence in cotton. Plant Physiol. 193(2):1177–1196.

119. Zhao F, Wang Z, Zhang Q, Zhao Y, Zhang H. 2006. Analysis of the physiological mechanism of salt-tolerant transgenic rice carrying a vacuolar Na+/H+ antiporter gene from Suaeda salsa. Journal of Plant Research 119, 95–104.

120. Zheng X, Jehanzeb M, Habiba, Zhang Y, Li L, Miao Y. 2019. Characterization of S40-like proteins and their roles in response to environmental cues and leaf senescence in rice. BMC Plant Biology 19.

121. Zhou X, Jiang Y, Yu D. 2011. WRKY22 transcription factor mediates dark-induced leaf senescence in Arabidopsis. Molecules and Cells 31, 303–313.

122. Zörb C, Noll A, Karl S, Leib K, Yan F, Schubert S. 2005. Molecular characterization of Na+/H+ antiporters (ZmNHX) of maize (Zea mays L.) and their expression under salt stress. Journal of Plant Physiology 162, 55–66.

